# Characterizing RNA stability genome-wide through combined analysis of PRO-seq and RNA-seq data

**DOI:** 10.1101/690644

**Authors:** Amit Blumberg, Yixin Zhao, Yi-Fei Huang, Noah Dukler, Edward J. Rice, Alexandra G. Chivu, Katie Krumholz, Charles G. Danko, Adam Siepel

## Abstract

The rate at which RNA molecules decay is a key determinant of cellular RNA concentrations, yet current approaches for measuring RNA half-lives are generally labor-intensive, limited in sensitivity, and/or disruptive to normal cellular processes. Here we introduce a simple method for estimating relative RNA half-lives that is based on two standard and widely available high-throughput assays: Precision Run-On and sequencing (PRO-seq) and RNA sequencing (RNA-seq). Our method treats PRO-seq as a measure of transcription rate and RNA-seq as a measure of RNA concentration, and estimates the rate of RNA decay required for a steady-state equilibrium. We show that this approach can be used to assay relative RNA half-lives genome-wide, with good accuracy and sensitivity for both coding and noncoding transcription units. Using a structural equation model (SEM), we test several features of transcription units, nearby DNA sequences, and nearby epigenomic marks for associations with RNA stability after controlling for their effects on transcription. We find that RNA splicing-related features are positively correlated with RNA stability, whereas features related to miRNA binding, DNA methylation, and G+C-richness are negatively correlated with RNA stability. Furthermore, we find that a measure based on U1-binding and polyadenylation sites distinguishes between unstable noncoding and stable coding transcripts but is not predictive of relative stability within the mRNA or lincRNA classes. We also identify several histone modifications that are associated with RNA stability. Together, our estimation method and systematic analysis shed light on the pervasive impacts of RNA stability on cellular RNA concentrations.

## Introduction

Gene regulation is an exquisitely complex process that operates at all stages of gene expression, ranging from pre-transcriptional chromatin remodeling to post-translational modification of proteins. However, the concentration of RNA molecules in the cell appears to serve as the primary target of many regulatory mechanisms. Many studies of gene regulation focus on the production of RNA, often at the stages of transcriptional pre-initiation, initiation, or release from pausing into productive elongation. RNA concentrations, however, result from a dynamic equilibrium between the production of new RNA molecules and their decay (Hao and Baltimore 2009; Rabani et al. 2011, 2014; Schwanhausser et al. 2011; Sharova et al. 2009; Tani et al. 2012; Yang et al. 2003). Indeed, bulk differences in RNA concentrations across types of transcription units (TUs) often result from differences in RNA decay rates. For example, protein-coding mRNAs, on average, are relatively stable, whereas lincRNAs are less stable, and enhancer RNAs (eRNAs) and other short noncoding RNAs tend to be extremely unstable (Rabani et al. 2014; Schwalb et al. 2016; Tani et al. 2012; Mukherjee et al. 2017). Among protein-coding genes, mRNAs associated with housekeeping functions tend to be stable, whereas those associated with regulation of transcription and apoptosis tend to have much shorter half-lives, probably to enable RNA concentrations to change rapidly in response to changing conditions (Herzog et al. 2017; Lam et al. 2001; Schwanhausser et al. 2011; Tani et al. 2012; Yang et al. 2003). In some cases, RNA decay is accelerated by condition- or cell-type-specific expression of micro-RNAs or RNA-binding proteins (Gosline et al. 2016; Rabani et al. 2014).

Over several decades, investigators have developed numerous methods for measuring RNA decay rates or half-lives (Hynes and Phillips 1976; Kim and Warner 1983; Wada and Becskei 2017). A classical approach to this problem is to measure the decay in RNA abundance over time following inhibition of transcription, often using actinomycin D (Hao and Baltimore 2009; Raghavan et al. 2002; Yang et al. 2003). More recently, many studies have employed a strategy that is less disruptive to cellular physiology, based on metabolic labeling of RNA transcripts with modified nucleotides. In this approach, the relative proportions of labeled and unlabeled transcripts are quantified as they change over time, following an initial introduction or removal of labeled nucleotides (Tani et al. 2012; Wada and Becskei 2017). Today, metabolic labeling is most commonly accomplished using the nucleotide analog 4-thiouridine (4sU), which is rapidly taken up by animal cells and can be biotinylated for affinity purification (Dolken et al. 2008; Kenzelmann et al. 2007; Rabani et al. 2011, 2014; Schwalb et al. 2016; Windhager et al. 2012). Related methods use chemical conversion of 4sU nucleotide analogs to allow identification by sequencing and avoid the need for affinity purification (Herzog et al. 2017; Schofield et al. 2018). In most of these assays, sample preparation and sequencing must be performed in a time course, making the protocols labor-intensive and dependent on the availability of abundant and homogeneous sample material (typically a cell culture). Many of these methods also have limited sensitivity for low-abundance transcripts. Owing to a variety of limitations, estimates of RNA half-lives tend to vary considerably across assays, with median half-lives often differing by factors of 2-3 or more (Tani et al. 2012; Wada and Becskei 2017). As yet, there exists no general-purpose assay for RNA half-life that is as robust, sensitive, or versatile as RNA-seq (Alkallas et al. 2017; Gaidatzis et al. 2015; Gosline et al. 2016) is for measuring cellular RNA concentrations, or PRO-seq (Kwak et al. 2013) and NET-seq (Churchman and Weissman 2011) are for mapping engaged RNA polymerases.

Recently, it has been shown that changes to RNA half-lives can be identified in a simpler manner, by working directly from high-throughput RNA-seq data (Alkallas et al. 2017; Gaidatzis et al. 2015; Gosline et al. 2016; Zeisel et al. 2011). The essential idea behind these methods is to treat RNA-seq read counts obtained from introns as a surrogate for transcription rates, and read counts obtained from exons as a surrogate for RNA abundance. Changes in half-life are then inferred from changes to the ratio of these quantities, under the assumption of a steady-state equilibrium between RNA production and decay. This approach assumes intronic read counts are representative of pre-mRNA abundances, when in fact they may derive from a variety of sources, and it can require a correction for differences in RNA processing rates (Alkallas et al. 2017). Moreover, the dependency on intronic reads limits the method to intron-containing transcription units that are transcribed at relatively high levels. Nevertheless, this simple approach requires no time course, metabolic labeling, transcriptional inhibition, or indeed, any experimental innovation beyond standard RNA-seq, making it an inexpensive and effective strategy for identifying genes undergoing cell-type- or condition-specific decay (Alkallas et al. 2017; Gaidatzis et al. 2015; Gosline et al. 2016).

In this article, we show that this same general approach—but using a measure of nascent transcription based on PRO-seq rather than intronic RNA-seq reads—results in improved estimates of relative RNA half-life. Our approach requires only two standard and widely applicable experimental protocols—PRO-seq and RNA-seq. It applies to intron-less as well as intron-containing transcription units; it requires no correction for RNA-processing rates; it makes efficient use of the available sample material and can be extended to tissue samples using ChRO-seq (Chu et al. 2018); it is relatively nondisruptive to the biological processes under study; and it is sufficiently sensitive to assay TUs expressed at low levels, including many noncoding RNAs (see **Supplemental Table 1** for a summary of advantages). We show, through a series of analyses, that these combined RNA-seq and PRO-seq measurements are a powerful means for assaying RNA stability that can reveal possible determinants of RNA decay.

## Results

### Matched PRO-seq and RNA-seq measurements are generally well correlated but suggest reduced stability of noncoding RNAs

We first compared PRO-seq and RNA-seq measurements for various TUs from across the human genome, to assess the degree to which transcriptional activity, as assayed by PRO-seq, is predictive of steady-state RNA concentrations, as assayed by RNA-seq. To reduce technical noise, we collected new data of each type in multiple replicates (two for PRO-seq, four for RNA-seq), all from the same source of K562 cells, and pooled the replicates after verifying high concordance between them (**Supplemental Fig. 1**). When analyzing these data, we considered all annotated TUs in GENCODE (Frankish et al. 2019), dividing them into mRNA (*n*=15,255), lincRNA (*n*=2,348), antisense (*n*=2,134), and pseudogene (*n*=1,274) classes. We quantified expression by the total number of mapped reads in transcripts per million (TPM), a measure that normalizes by both library size and TU length, and discarded TUs with insufficient read counts from either assay. Notably, we excluded the first 500 bp downstream of the TSS and 500bp upstream of TES for PRO-seq to avoid a bias from promoter-proximal pausing and polymerase deceleration (Kwak et al. 2013) (see **Methods**).

We found that the PRO-seq and RNA-seq measurements were well correlated overall, with Pearson’s *r*=0.82 (**Fig. 1**), suggesting that transcription explains the majority of the variance in mRNA levels. A parallel analysis based on pooled intronic reads from the same RNA-seq libraries showed only a slightly higher correlation, with *r*=0.87 (**Supplemental Fig. 2**). At the same time, there were considerable differences in the degree of correlation across classes of TUs, ranging from a high of *r*=0.86 for protein-coding mRNAs to *r*=0.72 for lincRNAs, *r*=0.68 for antisense genes, and only *r*=0.59 for pseudogenes (**Fig. 1**). Similarly, the slopes of the lines of best fit on the log/log scatter plots decreased substantially (by roughly 50%) from mRNAs to noncoding RNAs and pseudogenes. We observed similar patterns for intron-containing and intron-less genes, but reduced values of *r* and slopes overall in intron-less genes (**Supplemental Figs. 3 & 4**). Together, these observations suggest that RNA decay rates have a more pronounced effect on steady-state RNA levels in noncoding RNAs and pseudogenes. These differences remain when TUs are matched by expression level (**Methods**; **Supplemental Fig. 5**) and when the HeLa cell type is evaluated instead (**Supplemental Fig. 6**).

**Figure 1.**
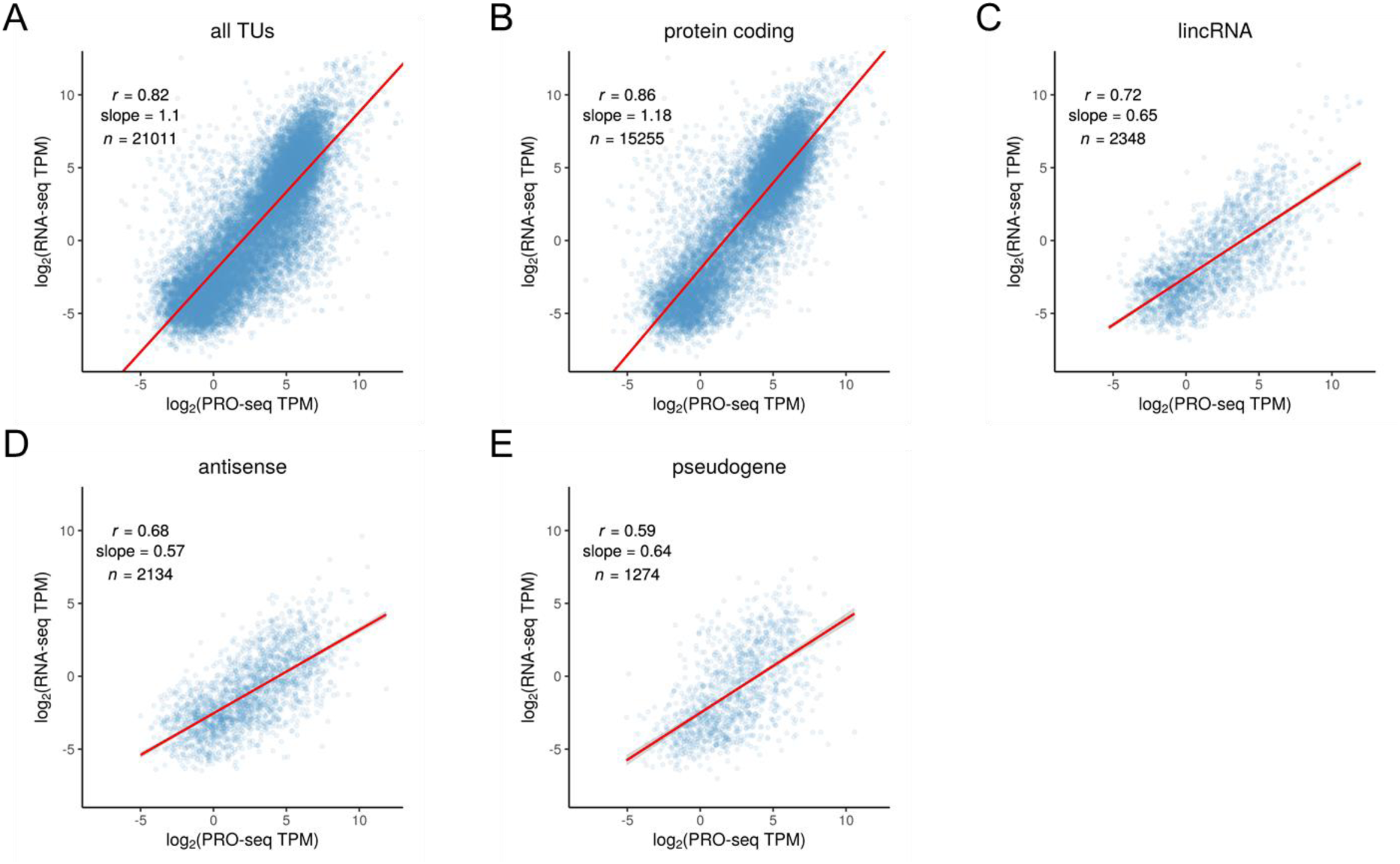
Scatter plots of PRO-seq vs. RNA-seq read counts for transcription units (TUs) in K562 cells, both shown in units of log_2_ transcripts per million (TPM) (see **Methods**). Panels describe (**A**) all annotated TUs (*n*=21,011), (**B**) protein-coding mRNAs (*n*=15,255), (**C**) intergenic lincRNAs (*n*=2,348), (**D**) intragenic antisense non-coding genes (*n*=2,134), and (**E**) pseudogenes (*n*=1,274), all from GENCODE (Frankish et al. 2019). For each plot, the linear regression line is shown together with Pearson’s correlation coefficient (*r*) and the slope of the regression line. Notice that as one proceeds from panel **B** to panel **E**, from mRNAs to noncoding RNAs and pseudogenes, there is a general decrease in both *r*, indicating greater variability of steady-state RNA concentrations at each transcription level, and the slope, indicating reduced average RNA concentrations for highly transcribed TUs.

Elongation rate is an important potential confounding factor in this analysis, because the PRO-seq density does not directly reflect the synthesis rate of RNA, but rather the synthesis rate divided by the elongation rate. However, when we correct for elongation rate using two different sets of estimates for K562 cells—one previously published (Veloso et al. 2014) and one based on our own experiments—we find that the correlation with RNA-seq measurements does not improve, and indeed, declines slightly. Thus the observed differences across classes of TUs do not appear to be driven primarily by differences in elongation rate (**Supplemental Text, Supplemental Fig. 7**, and **Discussion**).

### Relative RNA half-life can be estimated from the RNA-seq/PRO-seq ratio

As noted above, a quantity proportional to RNA half-life can be approximated in a straightforward manner from measurements of transcription rate and steady-state RNA concentration under equilibrium conditions (Alkallas et al. 2017; Gaidatzis et al. 2015). Briefly, if *β*_*i*_ is the rate of production of new RNAs for each TU *i, α*_*i*_ is the per-RNA-molecule rate of decay, and *M*_*i*_ is the number of RNA molecules, then, at steady state, *β*_*i*_ = *α*_*i*_ *M*_*i*_, and the decay rate can be estimated as *α*_*i*_ = *β*_*i*_ / *M*_*i*_ (see **Fig. 2A** & **Methods**). If we assume that *β*_*i*_ is approximately proportional to the normalized PRO-seq read counts for *i*, denoted *P*_*i*_, and *M*_*i*_ is proportional to the normalized RNA-seq read counts, denoted *R*_*i*_, then the ratio *P*_*i*_ / *R*_*i*_ is an estimator for a quantity proportional to the decay rate, and its inverse, *T*_1/2,*i*_^*PR*^ = *R*_*i*_ / *P*_*i*_, is an estimator for a quantity proportional to RNA half-life. As noted, the use of PRO-seq, rather than intronic read counts, for the measure of transcription has a number of advantages, including applicability to intron-less TUs and increased sensitivity for TUs expressed at low levels.

**Figure 2.**
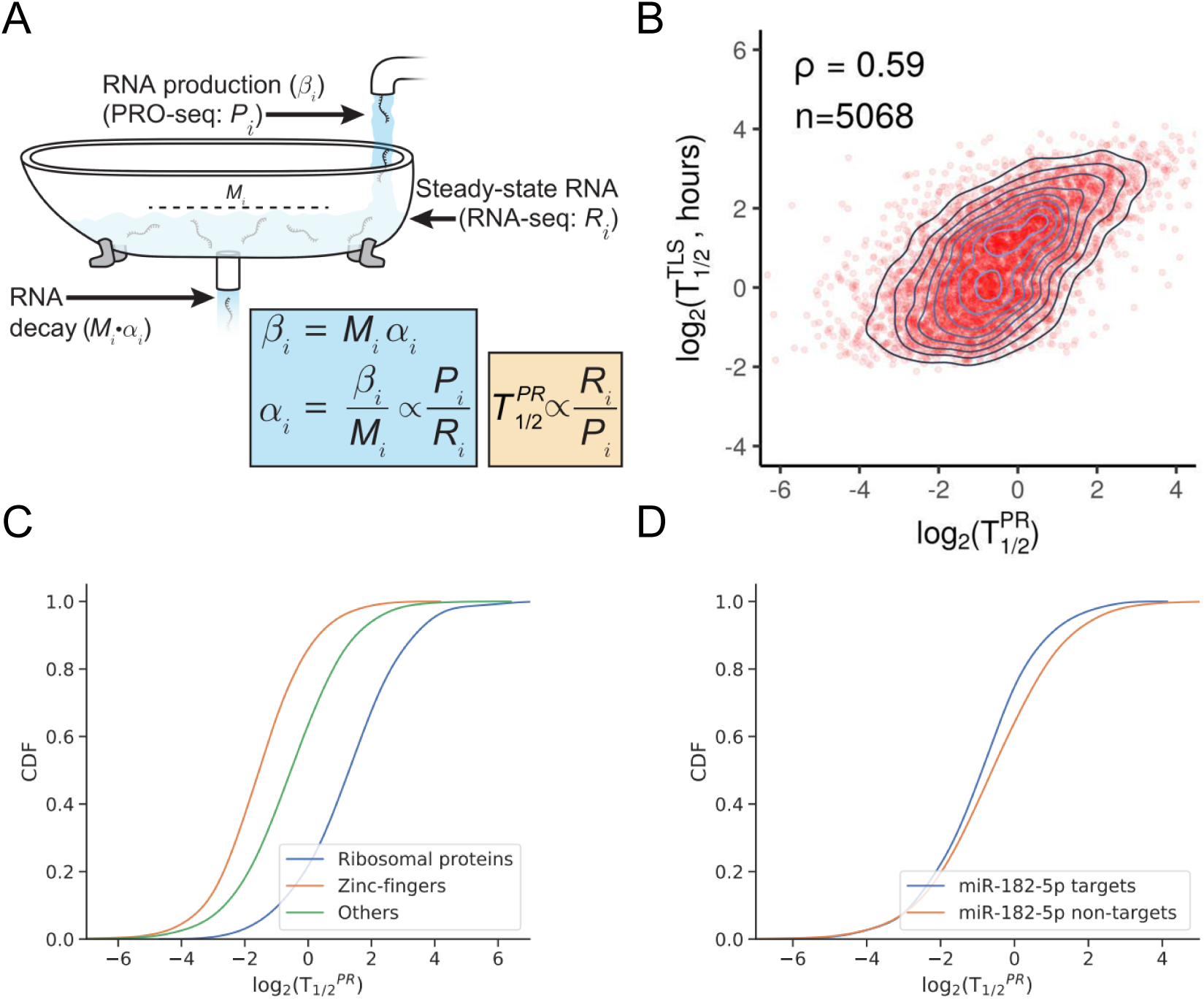
**(A)** Illustration of dynamic equilibrium between production and decay of RNA. PRO-seq (*P*_*i*_) can be used to measure production and RNA-seq (*R*_*i*_) to measure the resulting equilibrium RNA concentration. At steady-state, the production and decay rates must be equal, allowing for estimation of a quantity proportional to RNA half-life (*T*_1/2_^*PR*^) by the ratio *R*_*i*_ / *P*_*i*_ (see **Methods**). Illustration adapted from (Weingarten-Gabbay and Segal 2014). **(B)** Scatter plot with density contours for (log_2_) half-lives estimated by the PRO-seq/RNA-seq method (*T*_1/2_^*PR*^, *x*-axis) vs. those estimated by TimeLapse-seq (Schofield et al., 2018) (*T*_1/2_^*TLS*^, *y*-axis) for 5,068 TUs assayed by both methods in K562 cells. The *T*_1/2_^*PR*^ values are unit-less, whereas the *T*_1/2_^*TLS*^ values are expressed in hours. ρ = Spearman’s rank correlation coefficient. **(C)** Cumulative distribution functions (CDF) for (log_2_) estimated RNA half-lives, *T*_1/2_^*PR*^, for ribosomal proteins, zinc-finger proteins, and other genes (both comparisons have Kolmogorov–Smirnov test *p* = 3.99e-15). **(D)** Similar CDFs for mRNAs predicted to be targets of miR-182-5p vs. non-targets. K-S test *p* = 3.86e-10.

Following this approach, we estimated *T*_1/2_^*PR*^ values for TUs from across the genome using our PRO-seq and RNA-seq data for K562 cells. To validate our estimates, we compared them with estimates of RNA half-life for K562 cells from TimeLapse-seq (Schofield et al. 2018), a recently published method based on chemical conversion of 4sU. We compared our estimates of half-life with those from TimeLapse-seq (denoted *T*_1/2_^*TLS*^) at 5,068 genes measured by both methods. We found that the two sets of estimates were reasonably well correlated (Spearman’s ρ=0.59 **Fig. 2B**), especially considering the substantial differences in experimental protocols and the generally limited concordance of published half-life estimates across experimental methods (Tani et al. 2012; Wada and Becskei 2017). Moreover, if we remove the 50% of genes expressed at the lowest levels (as measured by PRO-seq), for which the noise contribution will tend to be largest, the correlation improves to ρ=0.62. By contrast, estimates based on intronic reads showed much poorer agreement with TimeLapse-seq (ρ=0.22; **Supplemental Fig. 8** and **Supplemental Text**), although it is worth noting that the correction for RNA processing introduced by Alkallas et al. (2017) could not be applied in our case, because it requires a comparison of two conditions. We found that our estimated *T*_1/2_^*PR*^ values were significantly shifted toward lower values for zinc finger proteins (**Fig. 2C**), many of which play key regulatory roles, and toward higher values for ribosomal proteins, which are representative of “housekeeping” genes. We also found that the predicted targets of numerous miRNAs, including the well-studied (Wei et al. 2015) miR-182 (**Fig. 2D**), have significantly reduced stability (see **Supplemental Fig. 9** for additional examples).

As further validation, we extended our comparison to include estimates of RNA half-life for K562 cells based on TT-seq (Wachutka et al. 2019), SLAM-seq (Wu et al. 2019), and the method of Mele et al. (2017), focusing on 3,991 genes for which estimates from all methods are available. In general, all methods show significant but somewhat modest levels of correlation in their half-life estimates, ranging from a high value of Spearman’s ρ=0.8 for the TimeLapse-seq and Mele et al. (2017) methods to a low of ρ=0.32 for SLAM-seq and our method (**Supplemental Fig. 10**). We attribute these differences in correlation to a variety of both technological and conceptual differences among methods (see **Discussion**). Notably, the TT-seq method—while similar to ours in some respects—has considerably reduced sensitivity, particularly for noncoding TUs (**Supplemental Tables 2 & 3**).

Finally, we explicitly adjusted our estimates of relative half-life for elongation rate, and found that the correlation with other methods did not improve (**Supplemental Figs. 11 & 12**). Moreover, we note that the variation across genes in estimated elongation rate is nearly an order of magnitude smaller than the variation in estimated half-lives, further indicating that elongation rate is not a dominant factor in our analysis, although it undoubtedly has some effect on our results (**Supplemental Material** and **Discussion**).

### Properties of transcription units that are predictive of RNA stability

To reveal potential determinants of RNA stability, we sought to identify features of TUs that were predictive of our estimated RNA half-lives. We focused on the mRNA and lincRNA classes, for which we could identify the most informative features. Anticipating an effect from splicing (Hamer and Leder 1979; Sharova et al. 2009), we focused our analysis on intron-containing TUs. We considered nine different features related to splicing patterns, transcript length, and G+C content (**Fig. 3** and **Supplemental Figs. 13 & 14**). In previous studies of this kind, investigators have examined the correlation of each feature with half-life, either individually or together in a multiple regression framework. By construction, however, *T*_1/2_^*PR*^ will tend to be statistically correlated with features predictive of transcription regardless of their true influence on half-life. Therefore, we instead made use of a Structural Equation Model (SEM) (Kaplan 2008) that explicitly describes the separate influences of features on transcription and half-life, and the contributions of both to RNA abundance (see **Methods** & **Fig. 3A**).

**Figure 3.**
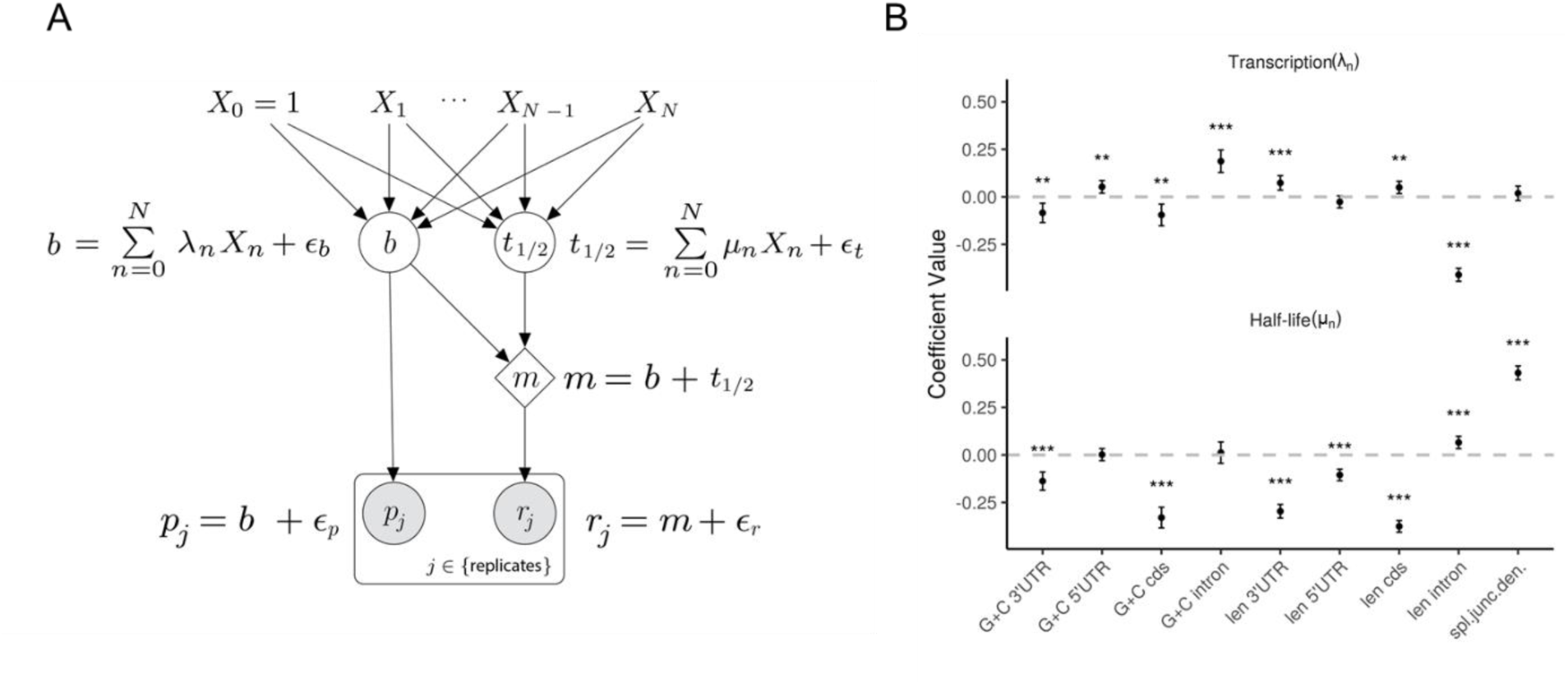
Features of transcription units (TUs) that are predictive of transcription rate and RNA half-life. **(A)** Structural Equation Model (SEM) describing the effects of an arbitrary collection of TU features (*X*_1_,…,*X*_*N*_, with intercept term *X*_0_=1) on transcription rate (*b*) and half-life (*t*_1/2_), as well as the downstream impact on mRNA concentration (*m*), normalized PRO-seq (*p*), and normalized RNA-seq (*r*) read counts. The model is linear in logarithmic space, with unmodeled variation accounted for as Gaussian noise (*ε*_*b*_, *ε*_*t*_, *ε*_*p*_, and *ε*_*r*_; see **Methods**). The coefficients for transcription rate (*λ*_*n*_) and half-life (*μ*_*n*_) are estimated by maximum likelihood, assuming independence of replicates and pooling data from all TUs of the same class. **(B)** Estimated values for coefficients for transcription (*λ*_*n*_; top) and half-life (*μ*_*n*_; bottom) for various features of interest. Results are for intron-containing mRNAs (see **Supplemental Figs. 13 & 14** for other classes). Features considered for each TU: G+C 3’UTR – GC content in 3’ UTR. G+C 5’UTR – GC content in 5’ UTR. G+C cds – GC content in coding region. G+C intron – GC content in intron(s). len 3’ UTR - length of 3’ UTR. len 5’ UTR - length of 5’ UTR. len cds – total length of coding region. len intron – total length of intron(s). spl. junc. dens. – number of splice junctions divided by mature RNA length. Error bars represent ±1.96 standard error, as calculated by the ‘lavaan’ R package (Yves 2012). Significance (from *Z*-score): * *p*<0.05; ** *p*<0.005; *** *p*<0.0005.

Our analysis revealed significant positive correlations with half-life of both splice junction density and total intron length, for intron-containing mRNAs and lincRNAs (**Fig. 3B; Supplemental Fig. 13**). The observation regarding splice junction density is consistent with previous reports for mRNAs (Hamer and Leder 1979; Sharova et al. 2009; Wang et al. 2002; Zhao and Hamilton 2007) and lincRNAs (Clark et al. 2012), as well as with the general tendency for intron-containing TUs to be more stable than intron-less TUs (**Supplemental Fig. 15**). The correlation with intron length is intriguing but could be an artefact of increased elongation rates in long introns (see below and **Discussion**). We also observed several patterns having to do with G+C content and length that are difficult to interpret owing to the complex correlations of these features with CpGs, transcription, splicing, and RNA half-life (see **Discussion and Supplemental Text**). In addition, we found that several features had coefficients of opposite sign for transcription and half-life (e.g., CDS, intron, and 3’UTR length), which could be driven, in part, by stabilizing selection on RNA levels (see **Discussion**).

To evaluate the degree to which these findings were influenced by elongation rate, we repeated the SEM analysis for the subset of 1,939 genes analyzed by Veloso et al. (2014), using an updated estimate of half-life that explicitly corrected for the estimated elongation rates of these genes (see **Supplemental Material**). We found that most of the results above held up under this analysis, with the main exception being the positive correlation between intron length and RNA half-life (**Supplementary Fig. 16**). This finding could indeed be an artifact of elongation rate in our uncorrected analysis because there is evidence of increased elongation rate (which would be perceived as reduced PRO-seq signal, and hence increased RNA-seq/PRO-seq ratio) in long introns (Gressel et al. 2017). We also observed some differences in the associations with G+C content.

As further validation, we performed a similar analysis using estimates of half-life based on TT-seq (Wachutka et al. 2019), SLAM-seq (Wu et al. 2019) and the study of Mele et al. (2017), focusing on 3,923 genes for which estimates were available from all methods (**Supplementary Fig. 17**). In these cases, we did not have separate measures of transcription and steady-state RNA abundance, so in place of the SEM analysis we performed multiple linear regression using the same features as covariates and the estimated half-lives from each of these other studies as outcomes. In general, the observed trends were similar across all methods. The major exceptions were intron length, where the other methods found a weak negative correlation instead of the positive correlation observed with our method, and 3’ UTR length, where the other methods found a weak positive correlation instead of a negative correlation. The intron length finding may again reflect a confounding influence from elongation rate. It is possible that the 3’ UTR length could similarly be influenced (in the other direction) by elongation rate, although in this case isoform selection may also play a role.

### DNA sequence correlates of RNA stability

Our estimates of RNA half-life for both coding and noncoding TUs provide an opportunity to better characterize DNA sequence correlates of RNA stability near transcription start sites (TSSs) (Almada et al. 2013; Core et al. 2014; Ntini et al. 2013; Sharova et al. 2009). We tested for associations between half-life and DNA words (*k*-mers) of various lengths near the TSS (**Supplemental Text**), but we found that the observed trends were predominantly driven by G+C content, with A+T-rich *k*-mers being enriched, and G+C-rich *k*-mers being depleted, in stable transcripts relative to unstable transcripts (**Fig. 4A**; **Supplemental Figs. 18–20**). Using the discriminative motif finder DREME (Bailey 2011), we identified several A+T-rich motifs associated with stable transcripts, and several G+C-rich motifs associated with unstable transcripts (**Fig. 4B&C**). Finally, we expanded our set of TUs to include previously identified eRNAs from K562 cells (Core et al. 2014) (see **Methods**), and found, interestingly, that stable eRNAs were slightly enriched, rather than depleted, for G+C-rich sequences (**Fig. 4A; Supplemental Fig. 20**). This trend was most strongly associated with CpG dinucleotides within 400bp of the TSS (**Supplemental Fig. 21**).

**Figure 4.**
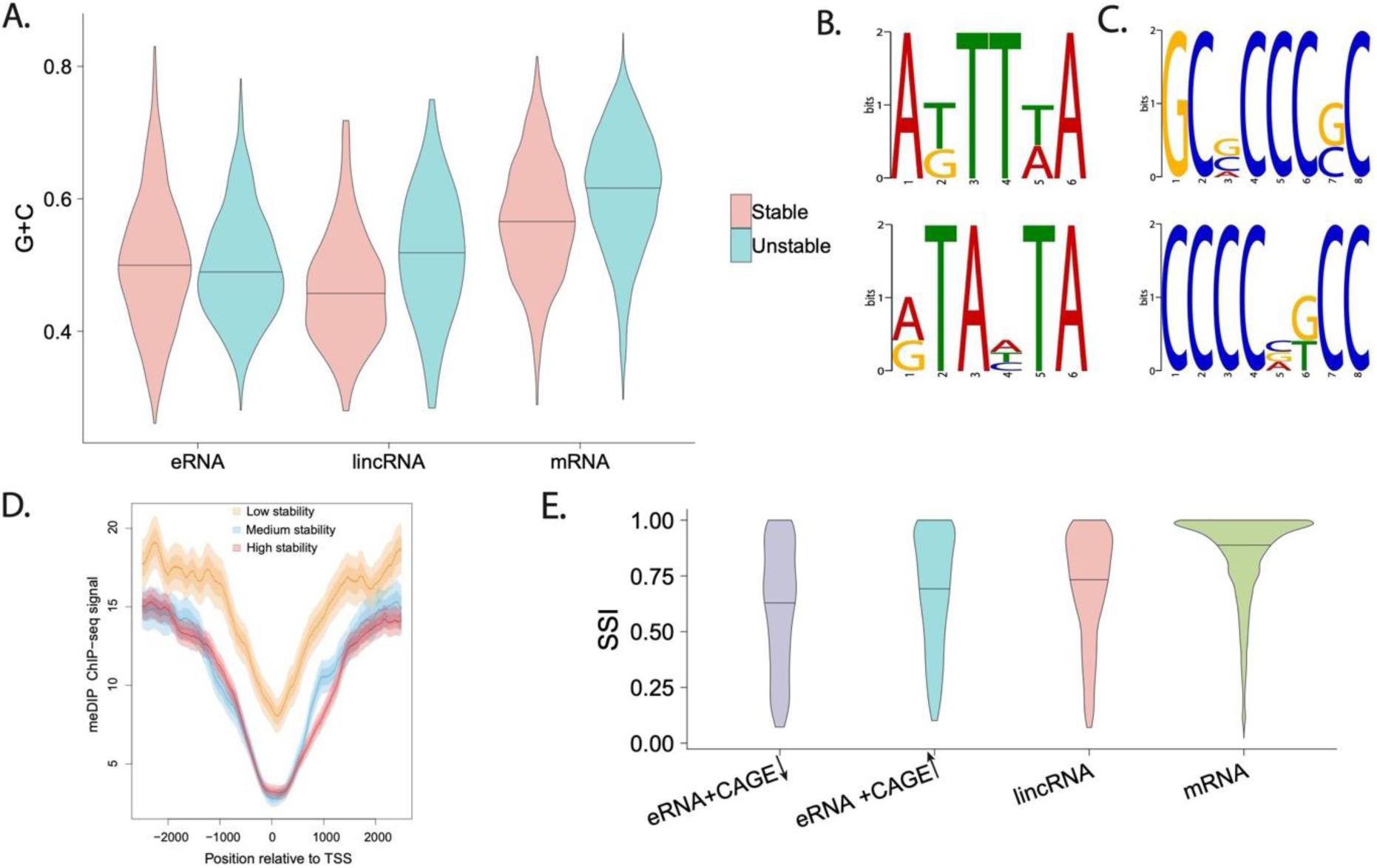
DNA-sequesnce, methylation, and RNA-binding-protein correlates of RNA stability near the TSS. **(A)** Distribution of G+C content (*y*-axis) for the 20% most (red) and least (blue) stable TUs, according to our estimated half-life (*T*_1/2_^*PR*^), in enhancer RNAs (eRNA), lincRNAs and mRNAs (*x*-axis). **(B&C)** Two most significantly enriched DNA sequence motifs in stable (B) and unstable (C) mRNAs. **(D)** Signal for MeDIP-measured DNA methylation for low-, medium-, and high-stability mRNAs (see **Methods**) as a function of distance from the TSS. Solid line represents mean signal and lighter shading represents standard error and 95% confidence interval. **(E)** Distribution of Sequence Stability Index (SSI) based on U1 and Polyadenylation sites (see **Methods**) for eRNAs, lincRNAs, and mRNAs. Separate plots are shown for eRNAs with low and high CAGE support, suggesting low and high stability, respectively.

The atypical patterns around CpG dinucleotides raise the possibility of an association with DNA methylation near the TSS. We therefore compared the methylation patterns of TUs exhibiting low, medium, or high levels of RNA stability, summarizing these patterns with meta-plots of average signal of the methylated DNA immunoprecipitation (MeDIP-seq) assay in K562 cells (ENCODE Project Consortium 2012; Vucic et al. 2009) as a function of distance from the TSS (**Supplemental Text**). We found that the medium- and high-stability TUs exhibited similar patterns of methylation, but the low-stability TUs show a clear enrichment (**Fig. 4D**). A similar trend was evident for lincRNAs (**Supplemental Fig. 22**). These observations suggest the possibility of epigenomic as well as DNA sequence differences associated with RNA stability, as we explore further below.

### U1 and Polyadenylation sites have limited predictive power for stability

We also directly tested for the possibility that differences in RNA half-life could reflect the presence or absence of either U1 binding sites (5’ splice sites) or polyadenylation sites (PAS) downstream of the TSS. Comparisons of (stable) protein-coding TUs and (unstable) upstream antisense RNA (uaRNA) TUs have revealed significant enrichments for proximal PAS in uaRNAs, suggesting that they may lead to early termination that triggers RNA decay. These studies have also found significant enrichments for U1 binding sites in protein-coding TUs, suggesting that splicing may play a role in enhancing RNA stability (Almada et al. 2013; Ntini et al. 2013). In previous work, we showed that these trends generalize to eRNAs as well. In particular, we found that a hidden Markov model (HMM) that distinguished between the occurrence of a PAS prior to a U1 site, and the occurrence of a U1 site prior to a PAS, could classify TUs as unstable or stable, respectively, with fairly high accuracy (Core et al. 2014).

We applied this HMM (see **Methods**) to our mRNA and lincRNA TUs and tested whether our DNA-sequence-based predictions of stability (as measured by a sequence stability index, or SSI) were predictive of our estimated *T*_1/2_^*PR*^ values. We also computed the SSI for the eRNAs identified from PRO-seq data and classified as stable or unstable based on CAGE data. We found that the mRNAs had the highest SSI, followed by lincRNAs, and then eRNAs (**Fig. 4E**), as expected. Interestingly, however, the subset of eRNAs that we find to be stable based on CAGE data also show elevated SSIs, roughly on par with lincRNAs. In addition, intron-containing lincRNAs have significantly higher SSIs than intron-less lincRNAs, although there was little difference in intron-containing and intron-less mRNAs (**Supplemental Fig. 23**). Moreover, within each of the mRNA and lincRNA groups, we found that the SSI changed relatively little as a function of *T*_1/2_^*PR*^, suggesting that the HMM had almost no predictive power for true RNA stability within these classes (**Supplemental Figs. 24 & 25**). These observations suggest that, whereas the U1 and PAS sequence signals do seem to distinguish broad classes of TUs with different levels of stability—namely, mRNAs, eRNAs, and uaRNAs—and the same signals are useful in distinguishing stable and unstable eRNAs, other factors likely dominate in determining gradations of stability within the mRNA and lincRNA classes (see **Discussion**).

### Additional epigenomic correlates of RNA stability

Finally, we asked whether other epigenomic marks such as histone modifications correlate with RNA stability. Histone modifications are primarily associated with transcriptional activity or repression, but they are also known to interact with splicing (Luco et al. 2010), and thus could influence RNA stability. Similar to the methylation analysis above (**Fig. 4D**), we produced meta-plots showing the average ChIP-seq signal in K562 cells as a function of distance from the TSS for 11 different common histone modifications (ENCODE Project Consortium 2012), separately for low-, medium-, and high-stability classes of expression-matched intron-containing mRNAs (see **Methods**). While some of these histone modifications did not differ substantially across stability classes, such as H3K9me1 and H3K9me3, several did show clear relationships with estimated RNA half-life (**Supplemental Fig. 26**). For example, H3k79me2, which is associated with transcriptional activity, gives a substantially higher signal in stable transcripts than in unstable ones, particularly in a peak about 1kb downstream from the TSS (**Fig. 5A)**. A similar pattern is observed for H3K4me2, H3K4me3, and H3K27ac. An inverse relationship is observed with H3K4me1, which is associated with active enhancers.

**Figure 5.**
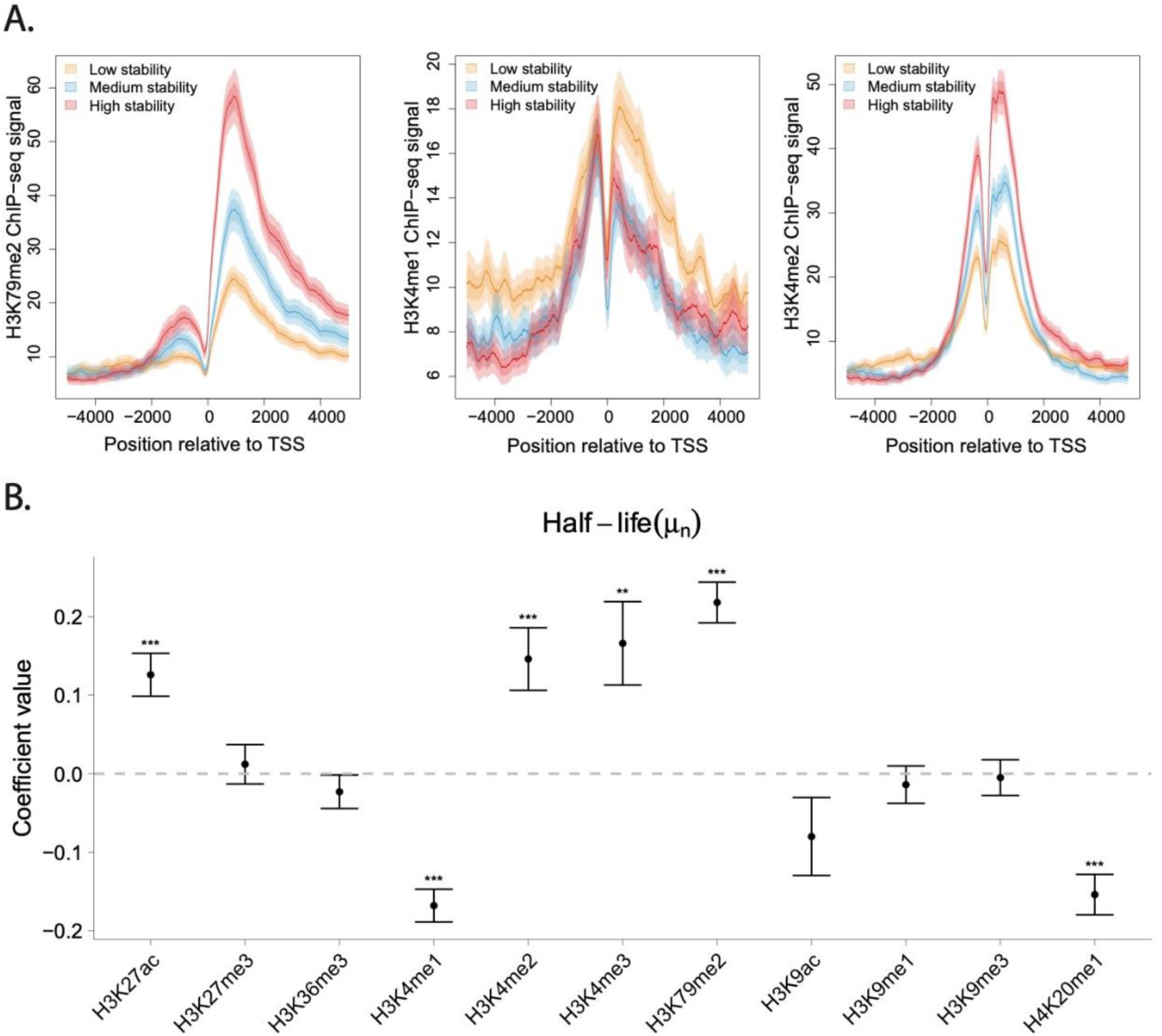
Histone-modification correlates of RNA stability. **(A)** ChIP-seq signal for H3K79me2 (*left*), H3K4me1 (*middle*), and H3K4me2 (*right*) for low-, medium-, and high-stability mRNAs (see **Methods**) as a function of distance from the TSS. Results are for intron-containing mRNAs matched by normalized PRO-seq signal. Solid line represents mean signal and lighter shading represents standard error and 95% confidence interval. **(B)** Estimated SEM coefficients for half-life (*μ*_*n*_) for 11 histone modifications, as assayed by ChIP-seq in the 500 bases immediately downstream of the TSS, also for intron-containing mRNAs (**Methods**; see **Supplemental Figs. 27-29** for additional results). Error bars and significance are as in **Fig. 3B**.

As an alternative strategy for identifying epigenomic correlates of RNA stability while correcting for transcription, we again applied our SEM framework, this time using the 11 histone marks as covariates for estimated RNA half-life and considering the ChIP-seq signals immediately downstream of each TSS (**Fig. 5B, Supplemental Fig. 27**). As expected, the strongest correlations were detected with transcription rate, and these generally had the expected sign, for example, with positive correlations for the activation marks H3K27ac, H3K4me1, H3K4me2, and H3K4me3, and negative correlations for the repressive marks H3K9me3 and H3K27me3. All of these patterns were consistent between lincRNAs and mRNAs (**Supplemental Fig. 27 & 28**), and they did not change substantially as a function of distance from the TSS (**Supplemental Fig. 29**). However, we did additionally identify several significant correlates of half-life. For mRNAs these were generally consistent with the ones identified from the ChIP-seq meta-plots, for example, with H3K79me2 showing a positive correlation with RNA half-life, and H3K4me1 showing a negative correlation. In general, the estimated coefficients were similar for mRNAs and lincRNAs, but there were some notable differences: for example, the activity mark H3K36me3, shows a strong negative correlation with RNA half-life in lincRNAs but a weaker and position-dependent positive or negative correlation with mRNA half-life; and the silencing marks H3K9me1 and H3K9me3 show positive correlations for lincRNA half-life but negative or near-zero correlations for mRNA half-life (**Supplemental Fig. 28**). These divergent patterns could possibly reflect differences in the degree to which splicing is co-transcriptional in mRNAs and lincRNAs (Tilgner et al. 2012).

## Discussion

In this article, we have introduced a simple method for estimating the RNA half-lives of TUs from across the genome based on matched RNA-seq and PRO-seq data sets. Like previous methods based on intronic reads, our method assumes equilibrium conditions and produces a relative measure of half-life only. Unlike these methods, however, the use of PRO-seq allows us to interrogate intron-less TUs and TUs that are expressed at low levels (e.g., **Supplemental Tables 2 & 3**). Moreover, even for intron-containing and abundantly expressed genes, the PRO-seq-based measurements appear to be considerably more accurate than those based on intronic reads. Our approach also has a number of advantages in comparison to existing methods for estimating RNA half-lives based on transcriptional inhibition or metabolic labeling. For example, it does not require collecting data in a time course, which enables efficient use of both time and sample material; it can make use of RNA-seq or PRO-seq data generated for other purposes; it is relatively nondisruptive of the biological processes under study; and it can be extended to tissue samples using ChRO-seq (Chu et al. 2018) (see **Supplemental Table 1**). We have shown that our measurements of relative half-life are useful in a wide variety of downstream analyses.

It is worth noting that our ability to assay noncoding RNAs derives in part from the use of RNA-seq data from total rRNA-depleted RNA rather than, say, oligoDT-enriched mature mRNA. As a consequence, our estimates of stability actually reflect a combination of RNA maturation steps, and likely underestimate the influence of RNA stability alone. In separate work (not shown), we recently analyzed K562 polyA+ RNA-seq data from the ENCODE project using our methods, and did observe a slight improvement in the correlation with other estimates of half-life. It is also worth noting that all of the available methods interrogate RNA stability for a particular cell type under a particular set of conditions. In most cases, it still remains unclear how RNA stability varies across conditions or cell-types.

In a comparison of half-life estimates from several methods that have all been applied to K562 cells, including TimeLapse-seq (Schofield et al., 2018), TT-seq (Wachutka et al. 2019), SLAM-seq (Wu et al. 2019), and the method of Mele et al. (2017), we found reasonable agreement across methods, but also substantial differences (**Supplemental Fig. 10**). Indeed, the average pairwise Pearson’s correlation coefficient between sets of estimates was only *ρ*=0.57. It is difficult at this stage to disentangle the sources of these differences. Most likely, they result both from experimental noise and from a combination of fundamental differences among methods, including whether the estimates are based on steady-state assumptions or time-course measurements, whether transcriptional inhibition or activation is used, how the rate of transcription is assayed, and whether RNA abundance is based on total RNA or polyA+ RNA. These differences may make some methods better for certain classes of TUs than others (e.g., coding vs. noncoding RNAs, lowly vs. highly expressed TUs, intron-containing vs. intron-less TUs, or RNAs that are or are not at equilibrium). More work will be required to clarify the relative strengths and weaknesses of the available methods.

One particularly important limitation of our method is that we use PRO-seq as a proxy for the rate of transcription, but in reality PRO-seq is a measure of the occupancy of engaged RNA polymerases, which reflects both the rate of transcription and the rate of elongation. The PRO-seq signal along a gene body is analogous to the headlight brightness on a highway at night; an increase in signal can reflect either an increased number of cars entering the highway (analogous to an increased rate of transcription), or a back-up in traffic (analogous to a decreased elongation rate). As a consequence, variation in *T*_1/2_^*PR*^ across TUs could in part be driven by variation in elongation rate. We attempted to control for this possibility in several ways. First, we explicitly corrected our estimates of transcription and half-life with estimates of elongation rate for the same cell type, using both previously published estimates (Veloso et al. 2014) and ones obtained through our own experiments (**Supplemental Material**). We found that the correction did not improve the correlation of PRO-seq and RNA-seq measurements (**Supplemental Fig. 7**), nor did it improve the agreement with independent estimates of half-life (**Supplemental Fig. 11**). Second, we repeated our analysis of features predictive of half-life with the corrected estimates and found that it did not substantially alter our results, with one notable exception (**Supplementary Figure 16**; discussed below). Third, we observed that the variation in elongation rate across genes is smaller by almost an order of magnitude than the variation in estimated half-lives, indicating that it can account for, at most, a small fraction of the observed variation (**Supplemental Text**). We conclude from these analyses that elongation rate does undoubtedly have some impact on our half-life estimates, but overall, the effects appear to be limited. However, more work will be needed to obtain more accurate and more comprehensive estimates of elongation rates, and to fully understand their impact on half-life estimates.

To identify features that are predictive of RNA half-life, we devised a structural equation model (SEM) that explicitly describes the separate effects of each feature on transcription and half-life, as well as the resulting impact on RNA concentrations, PRO-seq, and RNA-seq data. While multivariate regression has been used to identify features associated with RNA stability (Sharova et al. 2009), our analysis is the first, to our knowledge, to attempt to disentangle the separate influences of such features on transcription and RNA stability. It is worth noting that this framework could also be useful for estimators based on intronic reads. The results of the SEM analysis were consistent with previous findings in many respects, particularly regarding the association between RNA splicing and RNA stability. The mechanism underlying this relationship remains unclear, but it is known that the exon junction complex (EJC) remains bound to the mature mRNA after its transport to the cytoplasm and it has been proposed that EJC components may protect the mRNA from decay (Sharova et al. 2009; Zhao and Hamilton 2007). In addition to the previously reported positive correlation of splice junction density and RNA half-life, we also observed a positive correlation between intron length and half-life. This observation could potentially indicate that RNA stability is enhanced by recursive splice sites (Sibley et al. 2015) or extended contact with the spliceosome in long introns. However, we could not confirm this finding after our correction for elongation rate using a subset of our full gene set, and it may therefore be an artifact of increased elongation rates in long introns. More work will be needed to confirm or reject this association.

It has recently been reported that U1 binding sites are enriched, and polyadenylation sites are depleted, downstream of the TSS in stable mRNAs relative to unstable upstream antisense RNAs (uaRNAs) and enhancer RNAs (eRNAs), suggesting that RNA stability is determined, in part, by the DNA sequence near the TSS. In this study, we tested not only whether this “U1-PAS axis” could distinguish TUs in stable classes (mRNAs) from those in unstable classes (uaRNAs and eRNAs) but also how predictive it is of half-life within these classes. We confirmed that a U1-PAS-based “sequence stability index” (SSI) is generally elevated for mRNAs, intermediate for lincRNAs, and reduced for eRNAs. Furthermore, this SSI can distinguish between more and less stable eRNAs, as quantified using CAGE (**Fig. 4E**). Somewhat surprisingly, however, we found that the SSI has essentially no predictive power for relative RNA stability within the generally more stable mRNA and lincRNA classes (**Supplemental Figs. 24 & 25**). One possible explanation for this observation is that the U1-PAS axis determines a kind of early “checkpoint” for stable transcripts—for example, by ensuring that premature transcriptional termination is avoided—but that once a transcript has cleared this checkpoint, these DNA sequence features are no longer relevant in determining RNA stability. Instead, the relative stability of mRNAs and lincRNAs may be predominantly determined by splicing-related processes, binding by miRNAs or RBPs, or other posttranscriptional phenomena. More work will be needed to fully understand the mechanistic basis of these differences in stability.

Some of the associations that we observed with half-life concerned G+C content, but these observations are generally difficult to interpret. Indeed, even the comparatively straightforward question of the relationship between G+C content and transcriptional activity has a long and contradictory literature, with several studies finding correlations between them (Kudla et al. 2006; Urrutia and Hurst 2003; Versteeg et al. 2003), but others claiming that the relationship between G+C and transcription is weak, at best, once confounding factors such as genomic context are properly accounted for (Arhondakis et al. 2008; Sémon et al. 2005). Sharova et al. (2009) identified a fairly pronounced negative correlation between RNA stability and the prevalence of CpGs in the 5’UTR, which is not supported by our analysis—although we interrogated only G+C content, not CpGs, in the 5’UTR. These authors raised the intriguing hypothesis this correlation may reflect the activity of splicing-associated methyl CpG-binding proteins (Young et al. 2005), but, to our knowledge, this idea has not been tested experimentally. In any case, it seems unlikely that the complex relationships among G+C content, CpGs, transcription, RNA stability and downstream effects such as translational efficiency can be fully disentangled through post-hoc statistical analyses. Instead, this effort will require experiments that directly perturb individual features of interest and separately measure the effects on a variety of processes.

Our observations of epigenomic correlates of transcription and stability are similarly challenging to interpret. We identified several histone modifications that are significantly associated with increased or decreased half-life, but we cannot rule out the possibility that these correlations reflect indirect relationships with confounding variables not considered here. However, the effect is quite strong for certain marks (such as H3K79me2 and H3K4me2) and it is apparent both in direct comparisons of PRO-seq-matched TUs (**Fig. 5A**) and in the SEM setting (**Fig. 5B**). It therefore seems plausible that it has a direct mechanistic basis, perhaps involving factors that interact both with DNA-bound nucleosomes and the spliceosome. Divergent patterns for mRNAs and lincRNAs (**Supplemental Fig. 27**) suggest the possibility of differences in these splicing-associated processes. Additional work will be needed to test these hypotheses.

One general pattern that emerges from the SEM analysis of histone modifications is that the coefficients for transcription and half-life are often different from zero in opposite directions (**Supplemental Figs. 27-29**). This trend of anti-correlation was less prominent with the TU features, but we did observe it with CDS, intron, and 3’UTR length (**Fig. 3B**). A possible explanation for this pattern is that it is, at least in part, a reflection of stabilizing selection on gene expression. If selection tends to favor a particular RNA level for each TU, then mutations that increase transcription may tend to be compensated for by mutations that decrease RNA stability, and vice versa. Thus, stabilizing selection might result in a tendency for features that are positively correlated with one measure (transcription or stability) to be negatively correlated with the other. Notably, this type of hypothetical causal interrelationship between transcription and stability is not considered in our SEM, nor in any other statistical model of which we are aware. As a result, it may be difficult to distinguish correlations that have a direct, mechanistic basis (say, relating to transcription) from their indirect “echoes” (say, relating to half-life) resulting from evolutionary constraint. Despite this potential limitation, our framework remains useful for identifying potentially interesting correlations, whose mechanistic underpinnings can then be further investigated through direct experimental perturbation.

## Materials and Methods

### PRO-seq and RNA-seq data preparation and processing

To minimize technical differences, we sequenced new PRO-seq (*n*=2) and RNA-seq (*n*=4) libraries, generated from cells grown in the same flask under the same conditions. Human K562 cells were cultured using standard cell culture procedures and sterile techniques. The cells were cultured in RPMI-1640 media supplemented with 10% fetal bovine serum (FBS) and 1% penicillin/streptomycin. For PRO-seq, 3’ and 5’ adapters were ligated as described (Chu et al. 2018) followed by library preparation as previously published (Mahat et al. 2016). Sequencing was done by Novogene on a HiSeq instrument with paired-end reads of 2×150bp. For RNA-seq, RNA was extracted using the Trizol method (see https://assets.thermofisher.com/TFS-Assets/LSG/manuals/trizol_reagent.pdf), followed by rRNA depletion using the Ribozero HMR Gold kit. Libraries were prepared using the NEB kit with TruSeq RNAseq adaptors. Single-end sequencing (length=75) was performed on a NextSeq500 instrument by the RNA Sequencing Core at the College of Veterinary Medicine, Cornell University.

### Read mapping and transcript abundance estimation

Raw data files in fastq format were trimmed using Cutadapt (Martin 2011) with parameters (-j 0 -e 0.10 --minimum-length=10). Reads were then aligned using HISAT2 (Kim et al. 2019, 2) with default parameters (hisat2 --threads 4 -x {index} -U {input.reads} -S {output} --summary-file {log}). We used the GRCh38/hg38 reference genome and the associated GENCODE gene annotations. HTSeq (Anders et al. 2015) was used for read counting for RNA-seq and PRO-seq. Importantly, for the purposes of read counting with PRO-seq, we omitted the first 500 bases downstream of the TSS and 500 bases upstream of TES to avoid a bias in read counts from promoter proximal pausing and polymerase deceleration. Finally, we normalized read counts by converting them to transcripts per million (TPM) (Wagner et al. 2012) based on the length of each TU.

### Estimation of RNA half-life from RNA-seq and PRO-seq data

We assume a constant rate of production of new RNAs, *β*_*i*_, a constant per-RNA-molecular rate of decay, *α*_*i*_, and a number of RNA molecules, *M*_*i*_. At steady state, *β*_*i*_ = *α*_*i*_ *M*_*i*_; therefore the decay rate can be estimated as *α*_*i*_ = *β*_*i*_ / *M*_*i*_, and the half-life as *T*_1/2_ = ln(2) / *α*_*i*_ = ln(2) × *M*_*i*_ / *β*_*i*_. We further assume that the normalized PRO-seq read counts (omitting both regions near TSS and TES) are proportional to the rate of production of new RNAs, *P*_*i*_ ∝ *β*_*i*_, and that the normalized RNA-seq read counts are proportional to the number of RNA molecules, *R*_*i*_ ∝ *M*_*i*_. Therefore, *T*_1/2_ ∝ *R*_*i*_ / *P*_*i*_. We define our unit-less estimator of half-life as *T*_1/2_^*PR*^ = *R*_*i*_ / *P*_*i*_, where “*PR”* denotes a PRO-seq/RNA-seq-based estimator. Notice that these unit-less *T*_1/2_^*PR*^ values can be compared across experiments only up to a proportionality constant, unless the raw read counts have been appropriately normalized.

### Structural equation model (SEM)

To separate the effects of TU features on decay from the effects on transcription, we developed an SEM using the ‘lavaan’ R package (Yves 2012). Let *X*_*n*_ be the *n*-th feature associated with a TU. We assume that the logarithms of this TU’s transcription rate and half-life, i.e., *b* = log *β* and *t*_1/2_ = log *T*_1/2_^*PR*^, are linear combinations of the *X*_*n*_’s and a TU-level random effect: 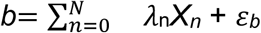 and 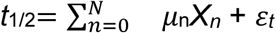 where *ϵ*_*b*_ ∼ *N*(0, *σ*_*b*_) and *ϵ*_*t*_ ∼ *N*(0, *σ*_*t*_) are independent Gaussian random variables explaining all variation not attributable to known features. Assuming a fixed value *X*_0_ = 1 for all genes, the parameters *λ*_0_ and *μ*_0_ can be interpreted as intercepts whereas *λ*_*n*≠0_ and *μ*_*n*≠0_ are regression coefficients indicating the contributions of feature *n* to transcription rate and half-life, respectively.

According to the model derived above, at steady state, *T*_1/2_^*PR*^∝ *M / β*, where *M* is the number of RNA molecules; therefore, *m* = log *M* is given by *m* = *b* + *t*_1/2_ + *C*, where *C* is an arbitrary constant that can be ignored here because it does not affect the estimation of regression coefficients. Denoting *p*_*j*_ = log *P*_*j*_ and *r*_*j*_ = log *R*_*j*_ as the logarithms of the PRO-seq and RNA-seq measurements in replicate *j*, respectively, we assume *p*_*j*_ ∼ *b* + *ε*_*p*_ and *r*_*j*_ ∼ *m* + *ε*_*r*_ where *ϵ*_*r*_ ∼ *N*(0, *σ*_*r*_) and *ϵ*_*r*_ ∼ *N*(0, *σ*_*r*_) are independent Gaussian random variables describing the noise in PRO-seq and RNA-seq experiments, respectively. Finally, we assume that all observations are independent across TUs. With these assumptions, and pooling information across TUs of the same class, we can estimate separate regression coefficients for transcription rates (*λ*_*n*_) and half-life (*μ*_*n*_) for all features by maximum likelihood.

### Transcription unit features

Transcription unit (TU) sequences were downloaded from BioMart using the R package biomaRt (Durinck et al. 2005, 2009). We considered only one isoform per annotated gene, i.e., selecting the longest transcript. Features based on properties of DNA sequences (e.g., G+C content) were then extracted using Biopython (Cock et al. 2009). The intron length was set equal to the transcript length minus the total exon length. The splice junction density was set equal to the intron number divided by the mature RNA length.

### eRNA analysis

We used eRNAs identified from our previous GRO-cap analysis in K562 cells (Core et al. 2014) restricting our analysis to putative eRNAs with divergent transcription (Danko et al. 2015) that fell at least 1kb away from annotated genes (*n*=21,816). To measure steady-state RNA levels, we used CAGE in place of RNA-seq owing to its greater sensitivity. We used the Nucleus PolyA and Non-polyA CAGE libraries from ENCODE. To measure transcription rates, we used PRO-seq data from same study (Core et al. 2014). For the stability analysis, we eliminated TUs having no mapped CAGE reads, and then selected the top 10% by CAGE/PRO-seq ratio as “stable” and the bottom 10% as “unstable”. These stable and unstable groups were then matched by PRO-seq signal (n=510).

### DNA word enrichments

We considered all DNA words (all possible combinations of A,C,G,T) of sizes *k* ∈ {2, 3, 4}. For each word *w*, we counted the total number of appearances in our set of stable TUs (top 20% by *T*_1/2_^*PR*^), denoted *c*_*s,w*_, and the total number of appearances in unstable TUs (bottom 20% by *T*_1/2_^*PR*^), denoted *c*_*u,w*_. These counts were collected in 1kb windows beginning at various distances downstream of the TSS (0, 500, 1000, and 1500 bp). The enrichment score for each word *w* and each window position was then computed as log_2_(*c*_*s,w*_/*c*_*u,w*_). A positive value of this score indicates an enrichment and a negative score indicates a depletion in stable TUs relative to unstable TUs. For eRNAs, we used a similar procedure but with 400 bp windows at distances of 0, 200, 400, and 600bp from the TSS.

### Motif discovery

For motif discovery, we used the discriminative motif finder ‘DREME’ (Bailey 2011) with default parameters (core width ranging from 3-7). For the stable motifs, we used the top 20% of TUs by *T*_1/2_^*PR*^ as the primary sequences and the bottom 20% as the control sequences. For the unstable motifs, we reversed the primary and control sequences.

### Sequence Stability Index (SSI)

We define the SSI to be the probability that a TU is “stable” based on our previously published U1-PAS hidden Markov model (HMM) (Core et al. 2014). Briefly, the HMM identifies a TU sequence as “stable” if either (1) it has a U1 splicing motif upstream of a PAS motif or (2) it lacks both a PAS motif and a U1 splicing motif, as detailed by Core et al. (2014). We applied the HMM to the first 1kb of sequence downstream of the annotated TSS and calculated the SSI as 1 minus the probability the TU is unstable, as output by the program. An implementation of the HMM is available at https://github.com/Danko-Lab/stabilityHMM.

### Matching by PRO-seq expression

We used the R package ‘MatchIt’ (Ho et al. 2007, 2011) to match groups of TUs by their normalized PRO-seq read counts (method=“nearset”). In cases of multiple groups, one group was selected as the reference and every other group was matched to that reference group.

### Metaplots

Metaplots showing the average values of signals of interest across loci (e.g., **Figs. 4D & 5A**) were produced using the ‘plotMeta’ function from the ‘Genomation’ (Akalin et al. 2015) R package. The input signal was provided in bigwig format and the loci were defined in bed format. In all cases, the average signal is plotted as a colored lined, with uncertainty indicated by the standard error of the mean (darker shading) and 95% confidence intervals (lighter shading) as specified by the “se” parameter.

### MicroRNA targets analysis

We obtained microRNA targets from TargetScanHuman (Agarwal et al. 2015), Release 7.2 (http://www.targetscan.org/vert_72/vert_72_data_download/Predicted_Targets_Info.def ault_predictions.txt.zip). We used all default predictions of conserved targets for each conserved miRNA family in the database.

### Gene categories

We obtained lists of genes encoding ribosomal proteins and zinc fingers from the HUGO Gene Nomenclature Committee (https://www.genenames.org/).

### Epigenomic Resources

Histone modifications, DNA methylation IP (MeDIP) and eCLIP data were downloaded from the ENCODE consortium (ENCODE Project Consortium 2012) as bigwig files annotated to the GRCh37/hg19 reference genome (https://www.encodeproject.org/).

### Software Availability

The software used for our data analysis and figure generation is available as Supplementary Material and via GitHub at https://github.com/EasyPiPi/blumberg_et_al.

## Acknowledgements

This research was supported, in part, by US National Institutes of Health grants R35-GM127070 (to AS) and R01-HG009309 (to CGD). The content is solely the responsibility of the authors and does not necessarily represent the official views of the US National Institutes of Health.

