## supplement for "Characterizing RNA stability genome-wide through combined analysis of PRO-seq and RNA-seq data"

### Supplemental Text

#### Comparison of half-life estimates with ones based on intronic reads

To compare our PRO-seq-based approach with an approach based on intronic reads, we repeated the estimation for about 21,000 TUs in our data set for which we could retrieve adequate numbers of RNA-seq reads mapped to introns (see **Methods**). We found that these intron-based estimates of half-life, $T_{1/2}^{IE}$, also correlated with the estimates from TimeLapse-seq but the correlation was substantially poorer than for the PRO-seq-based estimates (only $\rho$=0.22 for 3090 TUs accessible to both methods; **Supplemental Fig. 8**), suggesting that the PRO-seq-based approach provides less noisy estimates of transcription, and hence, reduced variance in estimates of half-life. Notably, however, the correction for RNA processing introduced by Alkallas et al. (2017) could not be applied in our case, because it requires a comparison of two conditions.

#### Correction for elongation rate in PRO-seq vs. RNA-seq correlation.

A potential confounding factor in the comparison of normalized read counts for PRO-seq and RNA-seq is elongation rate. Because PRO-seq read depth reflects a combination of transcription initiation rates and elongation rates (Danko et al. 2013; Jonkers et al. 2014), some reduction in correlation with RNA-seq could reflect variability across TUs in elongation rate. We investigated this possibility using two sets of measurements of elongation rate.

First, we made use of published estimates of elongation rate for the same K562 cell type (Veloso et al. 2014), focusing on ~2000 genes that overlap our set. We explicitly adjusted for the estimated elongation rates in our half-life calculation by multiplying them by the average PRO-seq abundance across gene bodies, under the assumption that the PRO-seq density is proportional to the synthesis rate divided by the elongation rate. After this adjustment, we observed no improvement (indeed, a slight decline) in the correlation of PRO-seq and RNA-seq measurements (**Supplemental Fig. 7**).

Second, we performed a new experiment to obtain our own estimates of elongation rate in K562 cells, and then corrected our half-life estimates for these measurements. To include as many genes as possible, we treated K562 cells with triptolide, which blocks the helicase activity of TFIIH and prevents new Pol II from initiating, and then carried out PRO-seq at time points of 0, 15, 30, 60, 120, and 240 min. We then estimated elongation rates for genes of sufficient length using the HMM-based methods we previously developed (Danko et al. 2013), slightly adapted to call receding rather than advancing polymerase "waves" (see below). These estimates turn out to have fairly high variance, for a combination of reasons, but they are generally consistent with those from Veloso et al. (2014) (**Supplemental Figure 12**), and when analyzed in bulk they give no indication that our estimates of relative half-life are dominated by differences in elongation rate (**Supplemental Figure 11**).

### Estimation of Elongation Rates Based on Triptolide Treatment

PRO-seq libraries were processed using proseq2.0 (https://github.com/Danko-Lab/proseq2.0), with parameters "-PE --UMI1=4 --UMI2=4". The bigwig files for each time point (0, 15, 30, 60, 120, and 240 min) were then used to infer a polymerase wave in the corresponding time interval (0 to 15 min, 0 to 30 min, etc.). A three-state Hidden Markov Model (HMM) (Danko et al. 2013) was used (https://github.com/Danko-Lab/polymeraseWaves). To allow for the receding rather than advancing polymerase waves, we subtracted the signal at each time point from the signal for the zero time point, rather than vice versa as in Danko et al (2013). We then used the differences between the two identified wave positions (e.g., those for 120 min and 60 min) divided by the corresponding time interval (e.g., 120 min - 60 min), to estimate the average elongation rate. We focused on estimates based on the 60 and 120 min time-points because these exhibited the lowest variance.

### Variation in Elongation Rate is Insufficient to Explain Variation in Half-Life

We additionally find that the variation in elongation rate across genes is smaller by almost an order of magnitude than the variation in estimated half-lives, indicating that it can account for, at most, a small fraction of the observed variation in half-life. In particular, the elongation rate estimates for the ~2000 genes from Veloso et al. (2014) range from 0.4 to 1.5 times the median value, if we discard the bottom and top 5% to avoid a strong influence from outliers. By contrast, our half-life estimates for the same genes range from 0.2 to 4.9 times the median value, again taking the 5th and 95th percentiles. Therefore, the half-life estimates exhibit a dynamic range of a factor of $4.9/0.2 \cong 25$, whereas the elongation rate estimates exhibit a dynamic range of a factor of only $1.5/0.4 \cong 3.75$.

### Validation of Features Predictive of Half-life

#### *Correction for Elongation Rate*

We redid our analysis of transcription unit (TU) features that are predictive of half-life after correcting for elongation rate, focusing on the ~2000 genes for which we had elongation rate estimates from Veloso et al. (2014). We found that most of the results reported in our manuscript held up under this analysis, with the important exception of the positive correlation between intron length and RNA half-life (**Supplementary Fig. 16;** see text for discussion). We also observed some differences in the associations with G+C content such as that G+C in introns was positively correlated with half-life and G+C in 5' UTRs was negatively correlated with half-life (both showed no significant correlation prior to the correction; **Supplementary Fig. 16**).

### DNA sequence correlates of RNA stability

We compared the sequences from 0 to 2500 bp downstream of the TSS for the 20% least and most stable TUs, according to the estimated $T_{1/2}^{PR}$, testing for global enrichments of all possible nucleotide *k*-mers in either class. We considered DNA word sizes of $k \in \{2, 3, 4\}$ and separately tested for enrichments in 1000 bp windows at various distances from the TSS (**Supplemental Figs. 18 & 19**). As above, we examined mRNAs and lincRNAs separately. In all cases, we matched the stable and unstable transcripts by their PRO-seq abundance estimates (see **Methods**) to avoid a bias from differences in transcription rates.

These tests identified a number of *k*-mers that were either enriched or depleted in stable transcripts, but these trends were almost completely explained by G+C content, with A+T-rich *k*-mers being enriched, and G+C-rich *k*-mers being depleted, in stable transcripts relative to unstable transcripts (**Fig. 4A**). Notice that, while this observation is generally consistent with the results of our SEM analysis, it specifically relates to G+C content near the TSS. Interestingly, these trends were largely independent of window position near the TSS, although they were slightly less pronounced in the first 500bp than in downstream windows **(Supplemental Fig. 20**). The patterns were similar for mRNAs and lincRNAs.

To shed further light on the local DNA sequence determinants of RNA stability, we expanded our set of TUs to include about 22,000 eRNAs from K562 cells, identified by GRO-cap in a previous study (Core et al. 2014). We obtained a relative measure of stability for these eRNAs based on the ratio of CAGE reads to PRO-seq reads within each TU (see **Methods**), using CAGE in this case because it was more sensitive than our RNA-seq data for eRNAs. We excluded eRNAs with no mapped CAGE reads. We then considered the 10% least stable and the 10% most stable of the remaining eRNAs ($n$=510 in each set), matching the two sets by normalized PRO-seq read counts downstream of the pause site. Interestingly, in this case, we found that stable eRNAs were enriched, rather than depleted, for G+C-rich sequences (**Fig. 4A**). This trend was strongly evident only for the first 400bp downstream of the TSS. It was especially pronounced for CpG dinucleotides (**Supplemental Fig. 21**). In contrast, AT (and to a lesser extent, TA) dinucleotides showed a fairly pronounced depletion for stable eRNAs.

### DNA methylation analysis

We partitioned our mRNAs, considering intron-containing TUs only, into five equally sized stability classes based on the estimated $T_{1/2}^{PR}$ values, and then subsampled from classes 1 (low stability), 3 (medium stability), and 5 (high stability) to obtain distributions matched by PRO-seq signal (see **Methods**). We then produced meta-plots for each of these three classes showing the average signal of the methylated DNA immunoprecipitation (MeDIP-seq) assay in K562 cells (Vucic et al. 2009; ENCODE Project Consortium 2012) as a function of distance from the TSS. We found that the medium- and high-stability TUs exhibited similar patterns of methylation, with intermediate levels 1-2kb upstream and downstream of the TSS, and a pronounced dip at the TSS which extended to about 1kb downstream (**Fig. 4D**). The low-stability TUs, by contrast, show a clearly distinct pattern, with elevated methylation levels across the whole region, no pronounced dip at the TSS, and a peak about 1kb downstream.

### Supplemental Table 1: Summary of Advantages of Method

| ***Advantage*** | ***Description*** |
|---|---|
| Efficiency of time and material | No need to collect data for full time course, reducing the number of experiments and the amounts of material needed |
| Reuse of existing data | RNA-seq and PRO-seq data sets can be decoupled. As a result, existing RNA-seq and/or PRO-seq data for adequately matched cells can be reused. |
| Other applications of data | Newly collected RNA-seq or PRO-seq data can be used for other purposes, e.g., analysis of proximal promoter pausing or identification of active enhancers (Danko et al. 2015) |
| Extension to tissue samples | Can be applied to tissue samples using ChRO-seq (Chu et al. 2018) |
| High sensitivity | Exploits high sensitivity of PRO-seq to noncoding and other low-abundance transcripts |
| Nondisruptive | Less disruptive to the biological processes under study than most drugs used for transcriptional inhibition and metabolic labeling. PRO-seq captures the positions of engaged RNA polymerases under the cellular conditions that exist when the experiment commences |
| Continual improvement | PRO-seq protocol is continually being improved. Current improvements allow PRO-seq libraries to be prepared in one day with comparable difficulty to enriching RNA-seq libraries for 4sU (Kim et al. 2020) |

### Supplemental Table 2: Numbers of annotated genes detected by PRO-seq and TT-seq

| | TPM>0 | | | TPM>1 | | |
|---|---|---|---|---|---|---|
| | **PRO-seq** | **TT-seq** | **TTseq/PROseq** | **PRO-seq** | **TT-seq** | **TTseq/PROseq** |
| protein_coding | 18496 | 17254 | 93.29% | 14359 | 11408 | 79.45% |
| lincRNA | 5829 | 4193 | 71.93% | 2725 | 1355 | 49.72% |
| antisense | 4578 | 3814 | 83.31% | 2862 | 1587 | 55.45% |
| pseudogene | 4982 | 3527 | 70.79% | 3094 | 1642 | 53.07% |

### Supplemental Table 3: Numbers of eRNAs detected by PRO-seq and TT-seq

| | **PRO-seq** | **TT-seq** | **TTseq/PROseq** |
|---|---|---|---|
| reads(TSS)>0 | 18639 | 13088 | 70.22% |
| reads(TSS)>1 | 15463 | 10866 | 70.27% |
| reads(TSS)>2 | 12923 | 9364 | 72.46% |

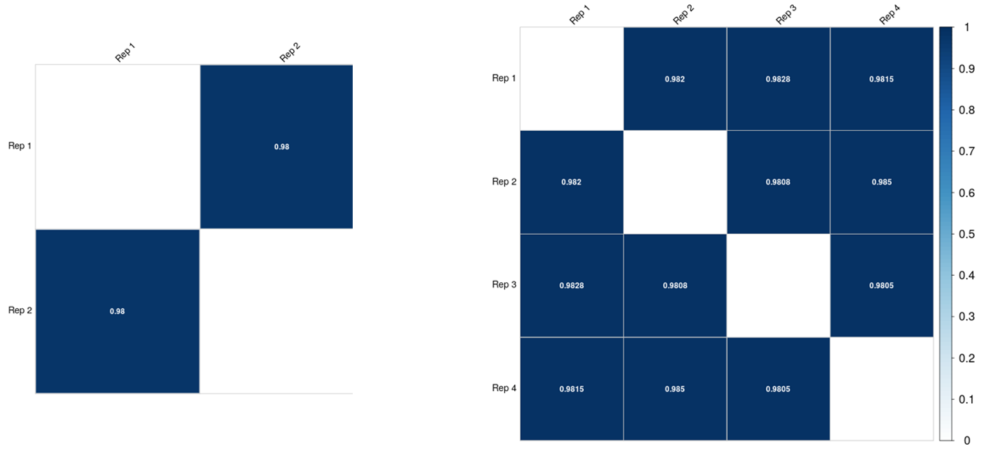


**Supplemental Figure 1. Correlation of PRO-seq and RNA-seq replicates in K562 cells.** (A) Pearson's correlation coefficient *r* for two replicates assayed by PRO-seq. (B) Pearson's correlations coefficient *r* for all pairs of four replicates assayed by RNA-seq. In both cases, genes without reads in one or more replicates were excluded.

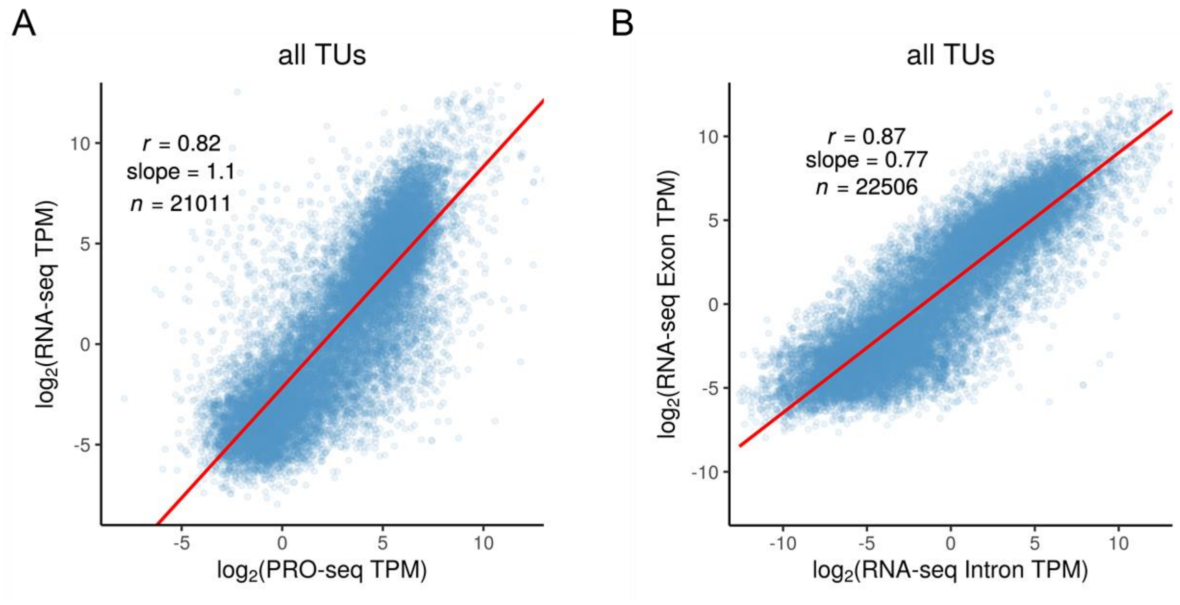


**Supplemental Figure 2. Correlation of exonic reads from RNA-seq with either PRO-seq or intronic reads.** (A) PRO-seq vs. exonic reads from RNA-seq. (B) Intronic reads vs. exonic reads from RNA-seq. Results are for all transcription units (TUs). *r* = Pearson's correlation coefficient.

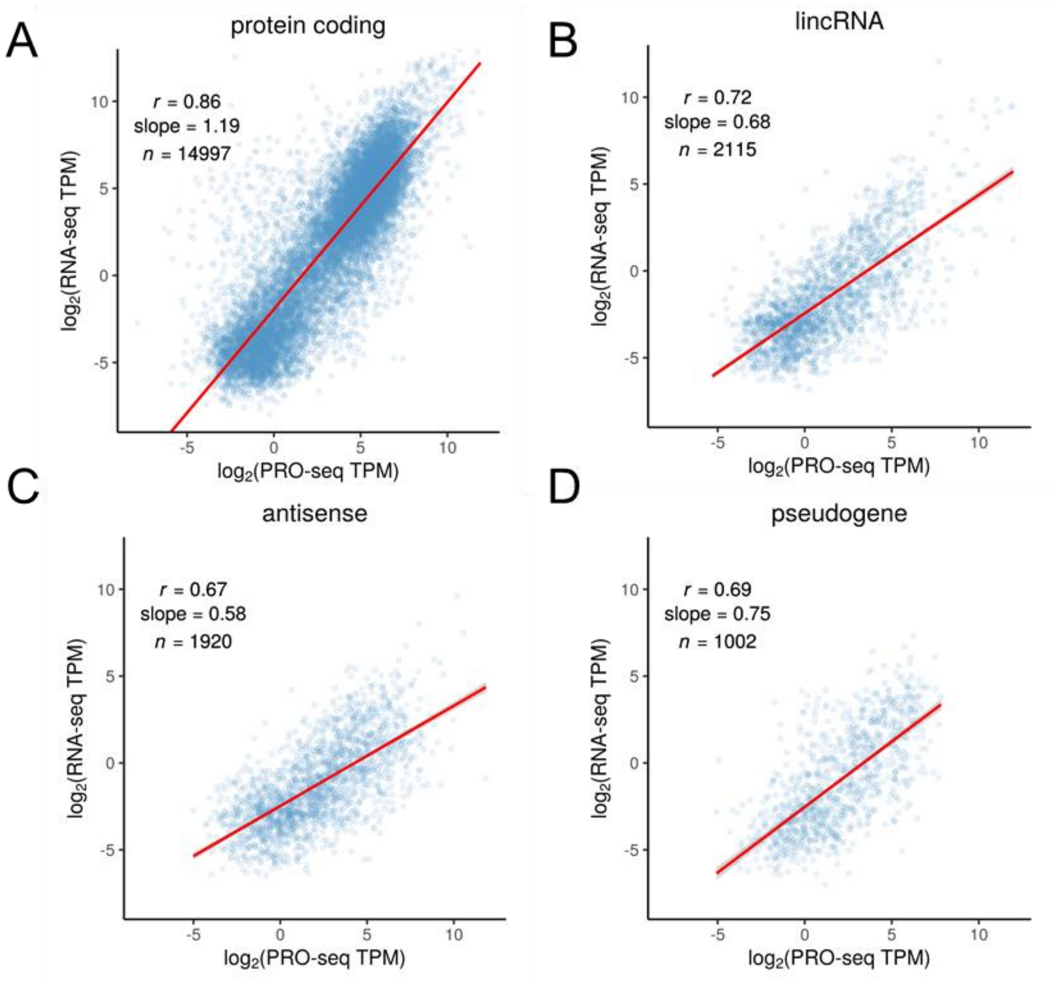


**Supplemental Figure 3. Scatter plots of PRO-seq vs. RNA-seq for intron-containing transcription units in K562 cells.** Panels describe (A) protein-coding mRNAs (B) intergenic lincRNAs (C) intragenic antisense non-coding genes, and (D) pseudogenes, all from GENCODE (Frankish et al. 2019). For each plot, the linear regression line is shown together with Pearson's correlation coefficient (*r*) and the slope of the regression line.

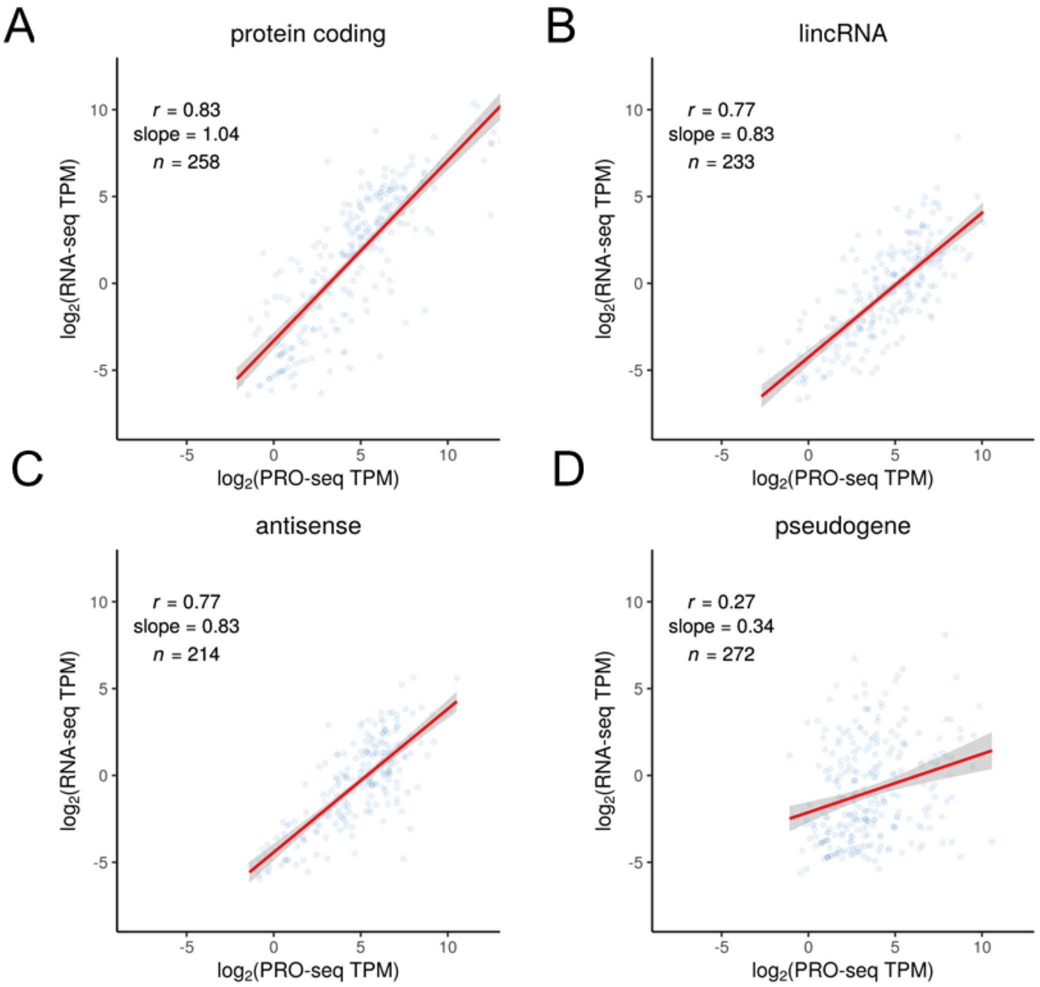


**Supplemental Figure 4. Scatter plots of PRO-seq vs. RNA-seq for intron-less transcription units in K562 cells.** Panels describe (A) protein-coding mRNAs (B) intergenic lincRNAs (C) intragenic antisense non-coding genes, and (D) pseudogenes, all from GENCODE (Frankish et al. 2019). For each plot, the linear regression line is shown together with Pearson's correlation coefficient (*r*) and the slope of the regression line.

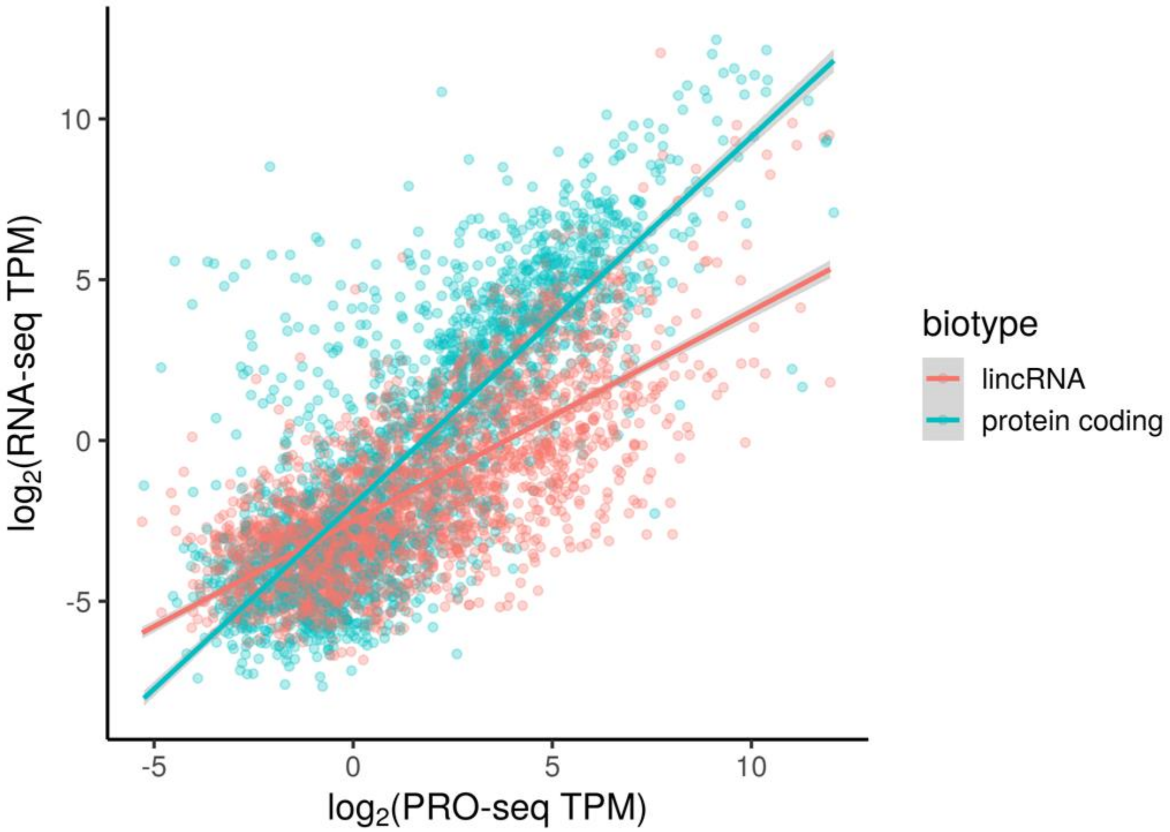


**Supplemental Figure 5. Scatter plots of PRO-seq vs. RNA-seq for lincRNAs and protein-coding mRNAs in K562 cells after matching by PRO-seq signal.** Subsampling was applied to obtain equal numbers of protein-coding mRNA and intergenic lincRNA (*n* = 2348) approximately matched by PRO-seq abundance (in TPM).

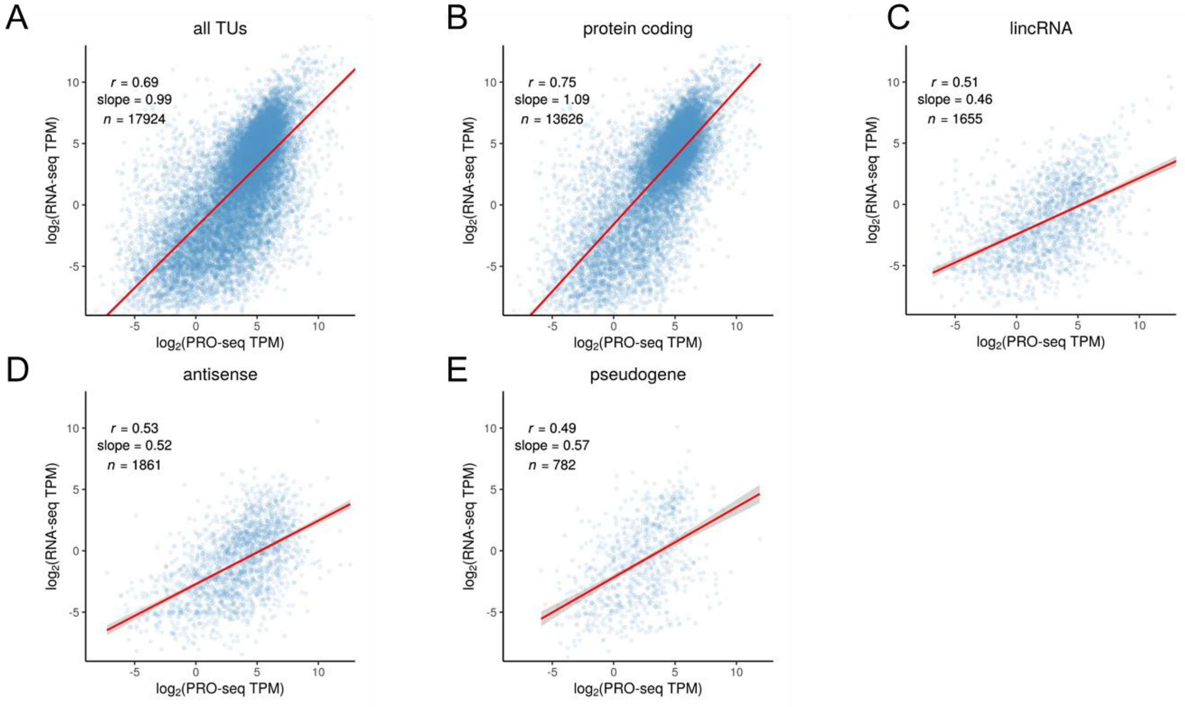


**Supplemental Figure 6. Scatter plots of PRO-seq vs. RNA-seq for intron-containing transcription units in HeLa cells.** Panels describe (A) all annotated TUs, (B) protein-coding mRNAs, (C) intergenic lincRNAs, (D) intragenic antisense non-coding genes, and (E) pseudogenes. For each plot, the linear regression line is shown together with Pearson's correlation coefficient (*r*) and the slope of the regression line.

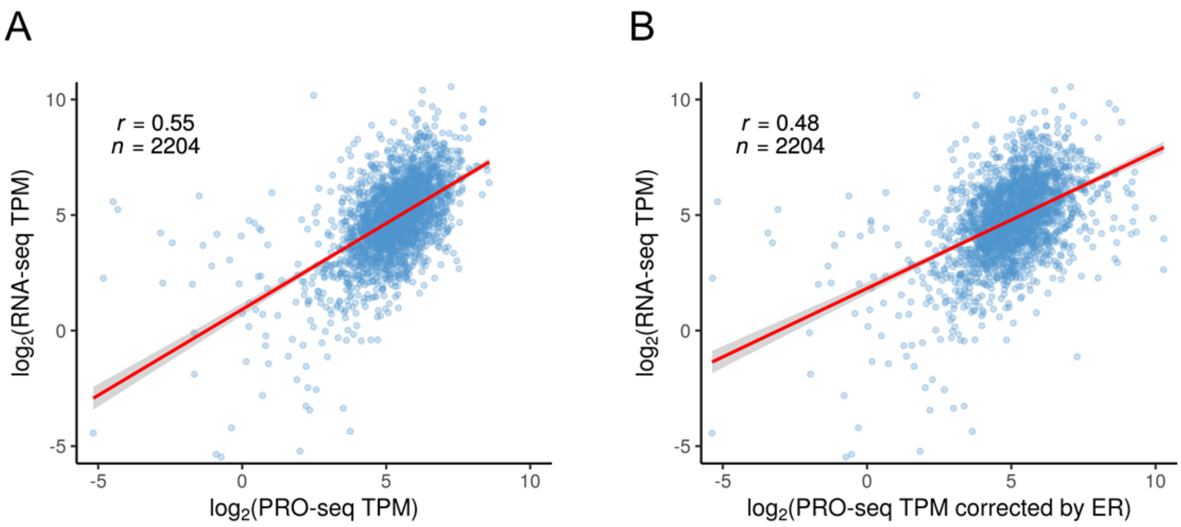


**Supplemental Figure 7. Scatter plots of PRO-seq vs. RNA-seq for lincRNAs and protein-coding mRNAs in K562 cells before and after correcting for elongation rate.** (A) With uncorrected PRO-seq estimates. (B) With corrected PRO-seq estimates. Both plots describe *n*=2204 genes for which estimates of elongation rate were available from Veloso et. al. (2014). Corrections in (B) were made by multiplying the PRO-seq TPM by the estimated elongation rate per gene (as described in the **Supplemental Text**). Similar results were obtained for an alternative set of elongation rate estimates based on our own experiments (see **Supplemental Text**).

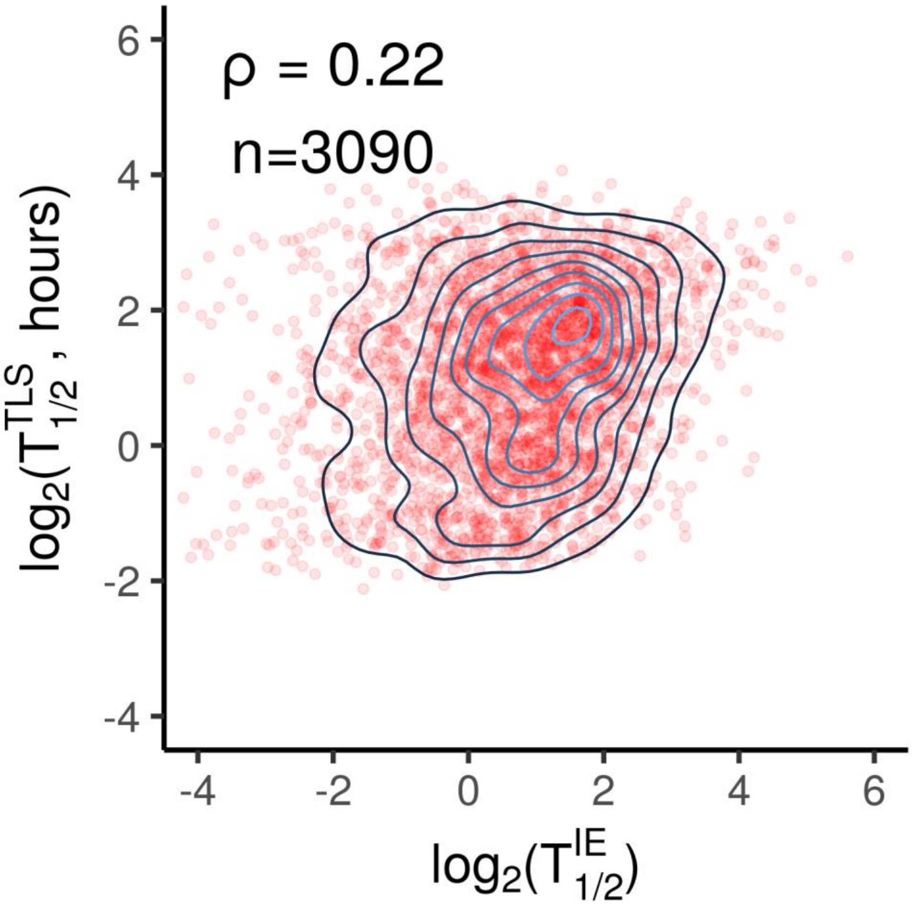


**Supplemental Figure 8. Intronic half-life vs. TimeLapse-seq half-life.** Scatter plot with density contours for $\log_2$(half-lives) estimated by the intronic/exonic RNA-seq method ($T_{1/2}^{IE}$, *x*-axis) vs. those estimated by TimeLapse-seq ($T_{1/2}^{TLS}$, *y*-axis) (*n* = 3090). ρ = Spearman's rank correlation coefficient.

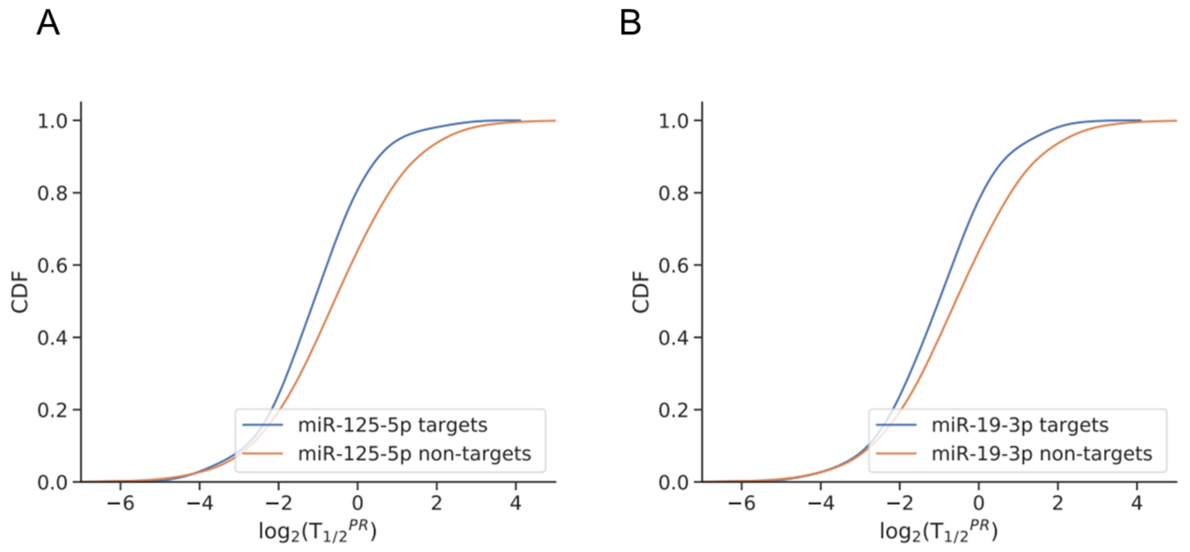


**Supplemental Figure 9.** Cumulative distribution functions (CDF) for estimated half-lives of predicted targets of miR-125-5p ($p$ = 4.30e-16, K-S test) and miR-19-3p ($p$ = 3.53e-15) vs. non-targets.

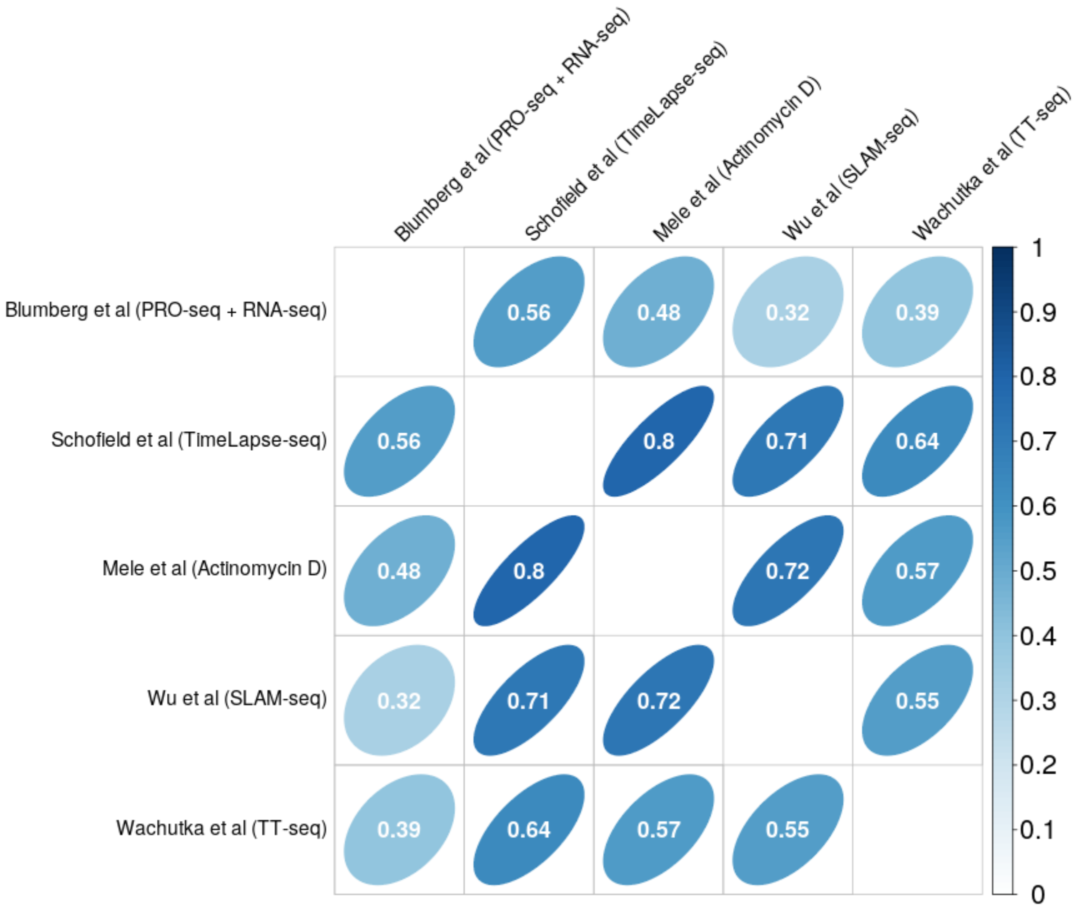


**Supplemental Figure 10. Correlation of estimated RNA half-lives under various methods.** Methods include our own, TimeLapse-seq (Schofield et al. 2018), the method of Mele et al. (2017), SLAM-seq (Wu et al. 2019), and TT-seq (Wachutka et al. 2019). All comparisons are based on a common set of protein-coding genes to which all methods had been applied ($n$ = 3991). The number in each cell of the matrix and the corresponding color represent Spearman's rank correlation coefficient ($\rho$) for each pairwise comparison.

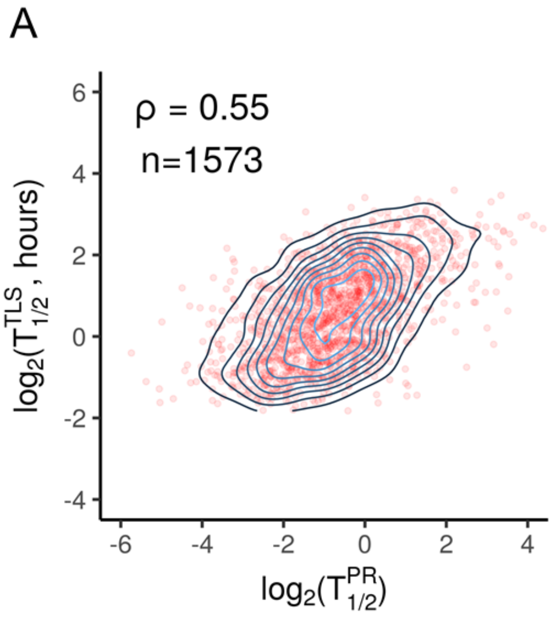


**Supplemental Figure 11. Correlation of PRO-seq-based half-lives ($T_{1/2}^{PR}$, *x*-axis) vs. estimates from TimeLapse-seq ($T_{1/2}^{TLS}$, *y*-axis) after correcting for elongation rate.** Elongation rates were obtained from (A) Veloso et. al. (2014). or (B) estimated in this study (see **Supplemental Text**). The correction was performed as described in **Supplementary Fig. 7** and the **Supplemental Text**. ρ = Spearman's rank correlation coefficient.

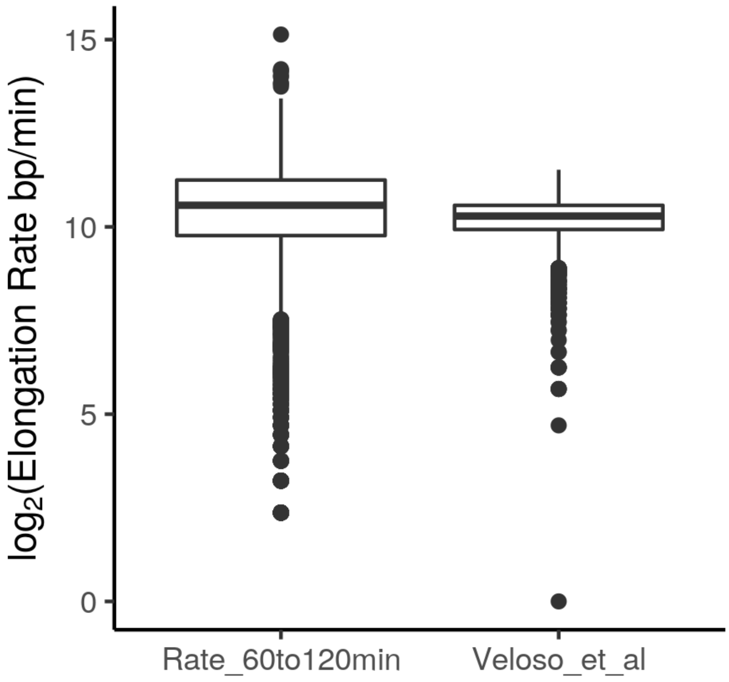


**Supplemental Figure 12. Distributions of Estimated Elongation Rates.** Elongation rates estimated from our own data (*left*; based on the 60 and 120 min time-points; see **Supplemental Text**) are compared with those from Veloso et. al. (2014) (*right*). Notice the similar median values (horizontal lines) but the larger variance in our estimates.

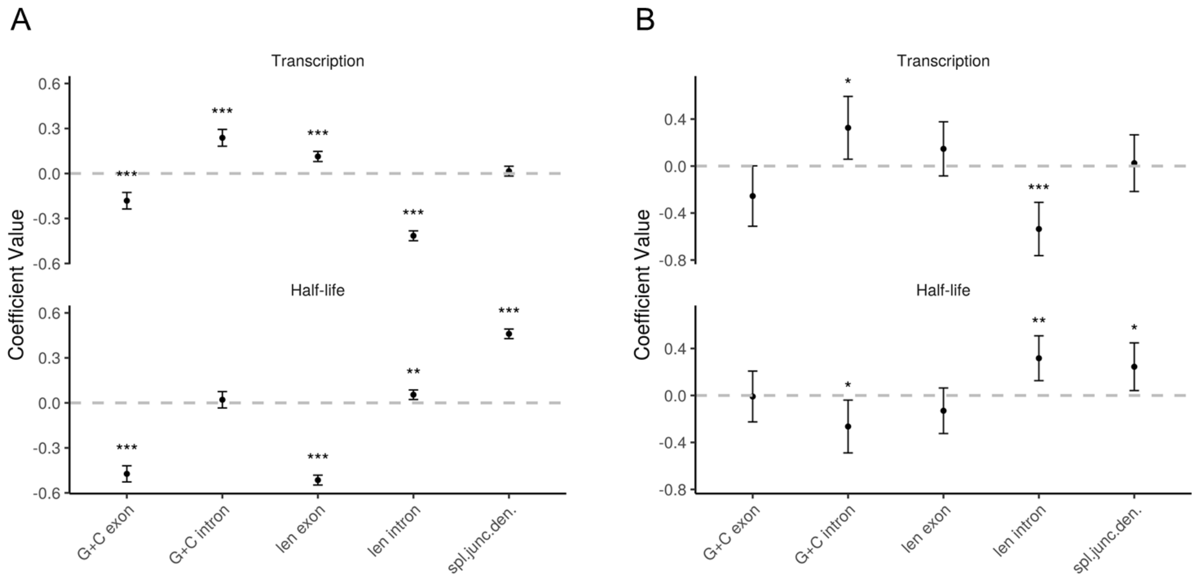


**Supplemental Figure 13. SEM results for features of intron-containing transcription units in K562 cells.** (A) Results for protein-coding mRNAs (*n* = 9685). (B) Results for lincRNA (*n* = 357). Features considered for each TU: G+C exon—GC content in exons; G+C Intron—G+C content in introns; len exon—Total exon length; len intron—Total intron length; spl. junc. den.—Number of splice junctions divided by mature RNA length. Error bars represent ±1.96 standard error, as calculated by the 'lavaan' R package (Yves, 2012). Significance (from *Z*-score): * $p<0.05$; ** $p<0.005$; *** $p<0.0005$.

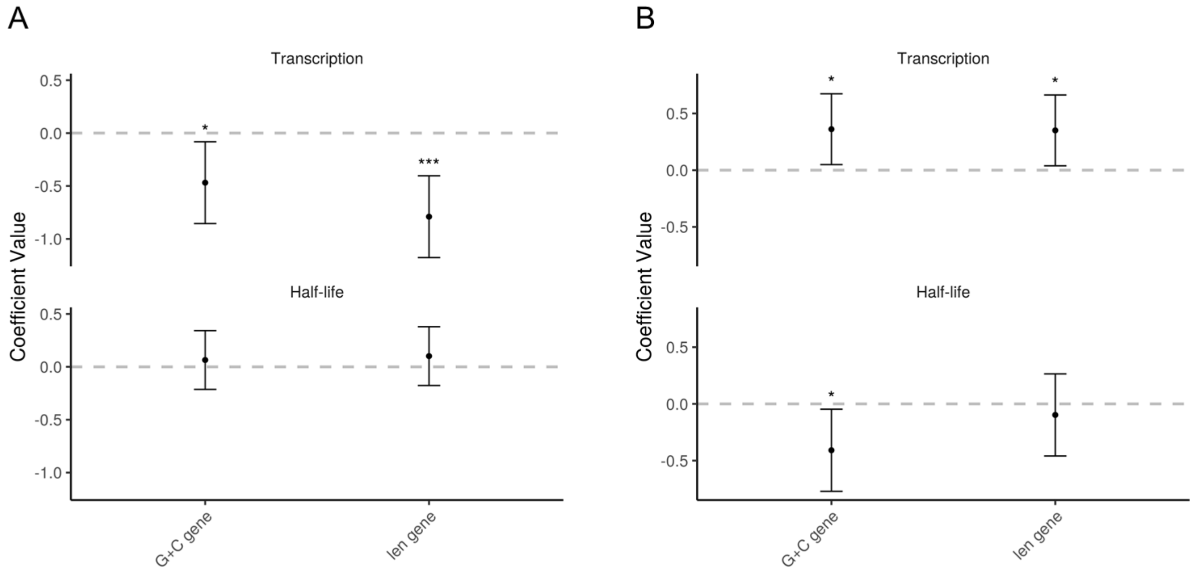


**Supplemental Figure 14. SEM results for features of intron-less transcripts in K562 cells.** (A) Results for protein-coding mRNAs (*n* = 145). (B) Results for lincRNAs (*n* = 77). Features considered for each TU: len gene—TU length; G+C gene—G+C content. Error bars represent ±1.96 standard error, as calculated by the 'lavaan' R package (Yves, 2012). Significance (from *Z*-score): * $p<0.05$; ** $p<0.005$; *** $p<0.0005$.

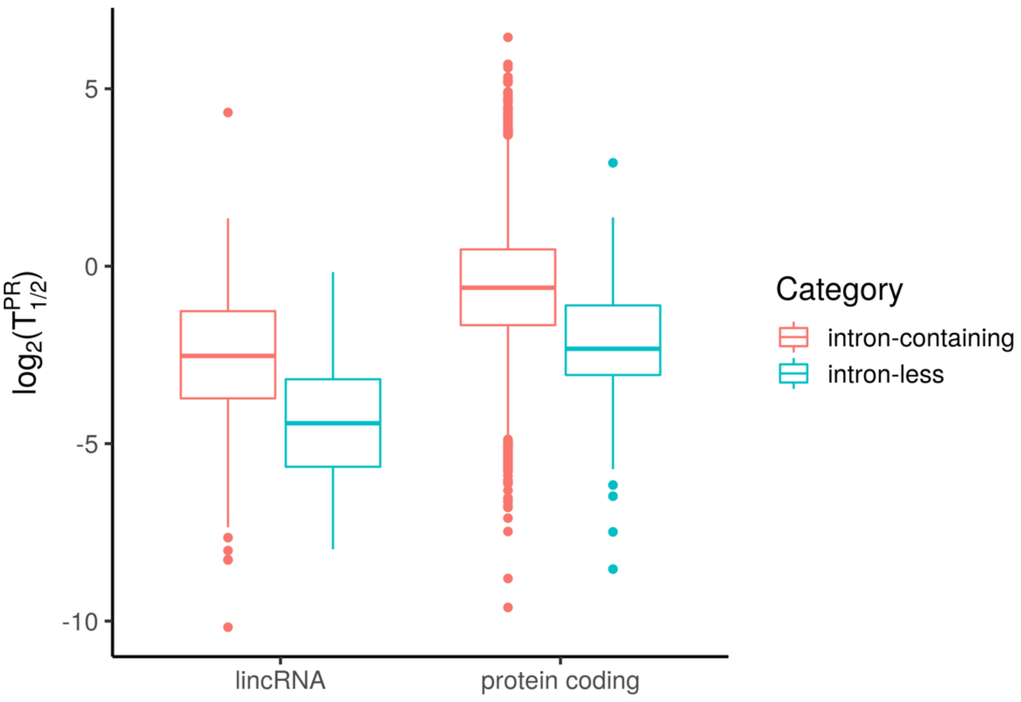


**Supplemental Figure 15. Estimated half-lives for intron-containing and intron-less transcription units.** Results for lincRNAs are shown on the *left* and results for protein-coding mRNAs are shown on the *right*.

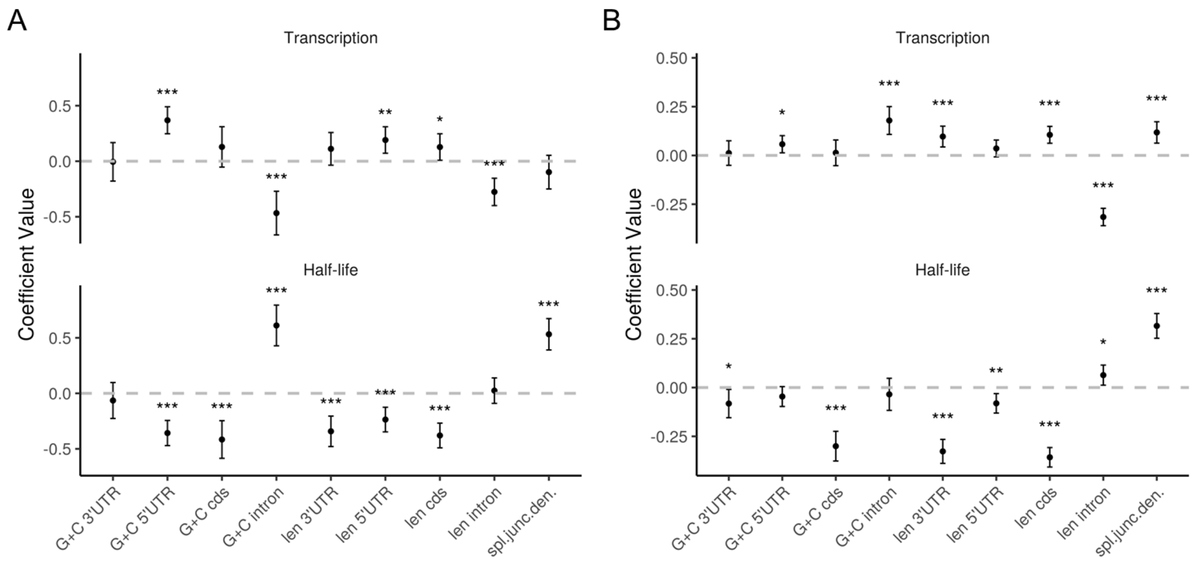


**Supplemental Figure 16. SEM results for features of intron-containing transcripts in K562 cells, with and without a correction for elongation rate.** (A) Results based on estimates of half-life that were explicitly corrected for elongation rates obtained from Veloso et al. (2014) as described in the **Supplemental Text**. (B) Results for the same set of genes without the correction for elongation rate. In both analyses, we used *n* = 1939 genes for which elongation-rate estimates were available.

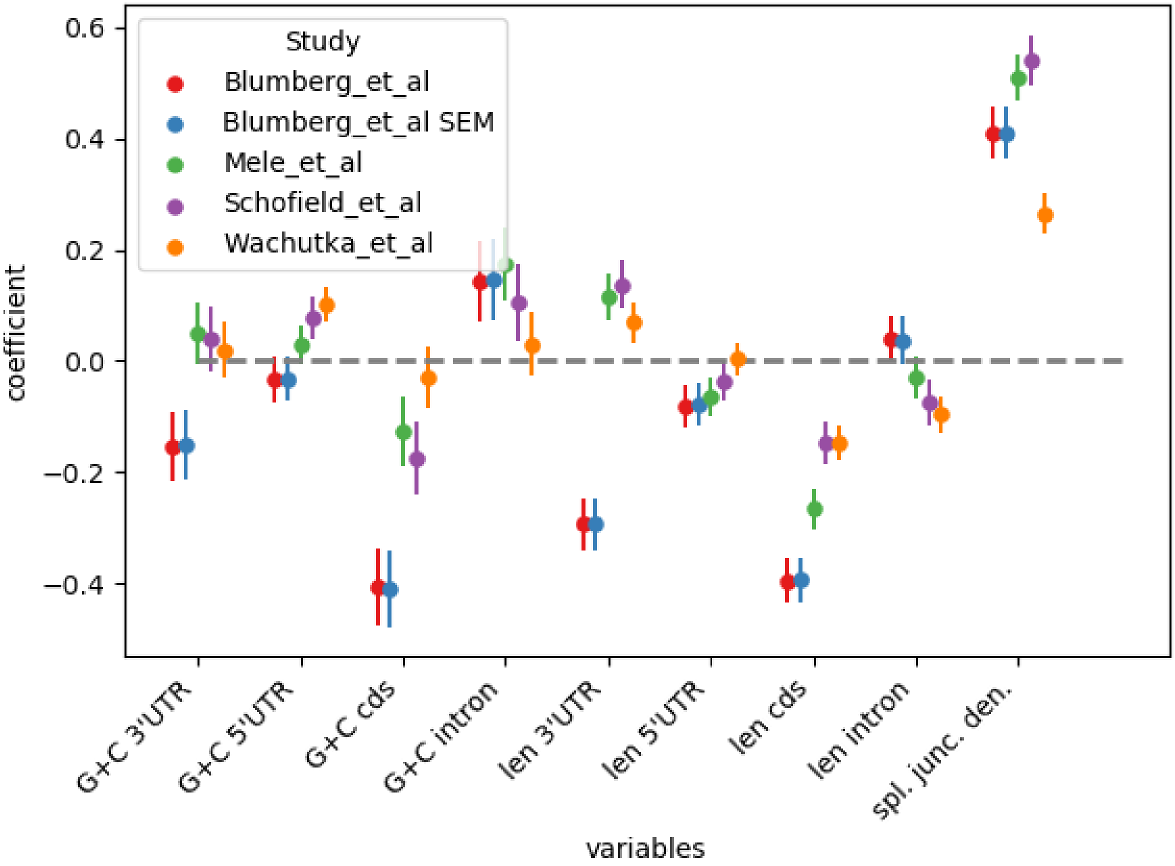


**Supplemental Figure 17. Multiple linear regression for features of transcription units versus RNA stability in K562 cells.** The same analysis was repeated with half-life estimates from our study (Blumberg et al.), the study of Mele et al. (2017), TimeLapse-seq (Schofield et al. 2018), and TT-seq (Wachutka et al. 2019). For comparison, we also show the coefficients from our SEM analysis (Blumberg et al. SEM). Results are based on *n*=3923 genes to which all methods had been applied.

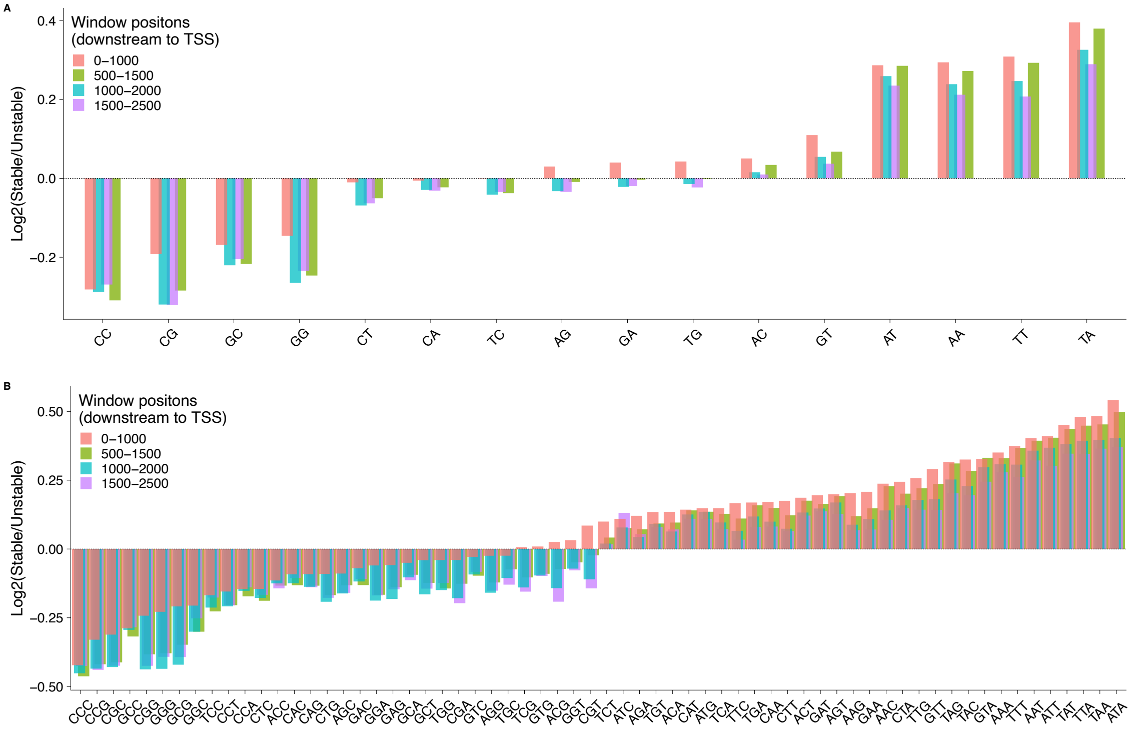


**Supplemental Figure 18. DNA word enrichments in stable transcripts for protein-coding mRNAs.** For each DNA word, we counted the numbers of total appearances of that word in all stable TUs (taking the top 20% by estimated half-life), $c_{s,w}$, and in all unstable transcripts (the bottom 20%), $c_{u,w}$. We then measured the stability enrichment as $\log_2(c_{s,w}/c_{u,w})$ (see **Methods**). The analysis was repeated in 1kb windows at four different distances downstream of the TSS (see legend). (A) Results for 2-mers. (B) Results for 3-mers. The order of the k-mers reflects the $(c_{s,w}/c_{u,w})$ ratio in the first window (0-1000bp).

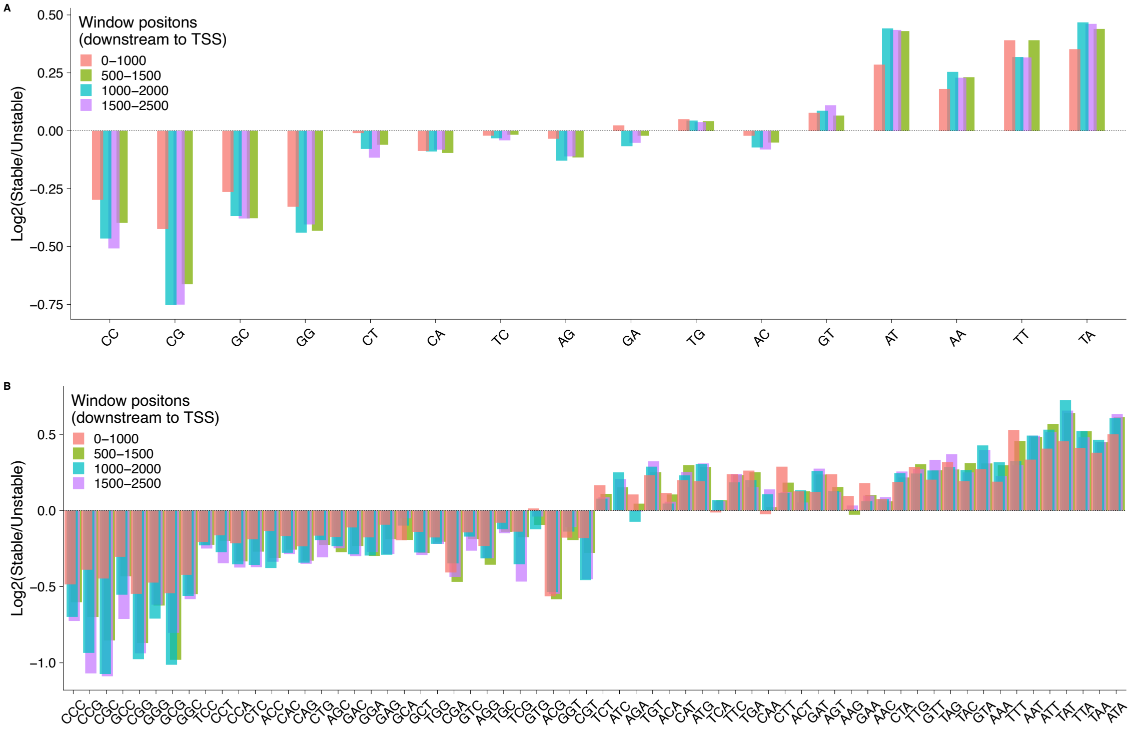


**Supplemental Figure 19. DNA word enrichments in stable transcripts for lincRNAs.** For each DNA word, we counted the numbers of total appearances of that word in all stable TUs (taking the top 20% by estimated half-life), $c_{s,w}$, and in all unstable transcripts (the bottom 20%), $c_{u,w}$. We then measured the stability enrichment as $\log_2(c_{s,w}/c_{u,w})$ (see **Methods**). The analysis was repeated in 1kb windows at four different distances downstream of the TSS (see legend). (A) Results for 2-mers. (B) Results for 3-mers. The order of the k-mers reflects the $(c_{s,w}/c_{u,w})$ ratio in the first mRNA window (0-1000bp; see **Supplemental Fig. 18**).

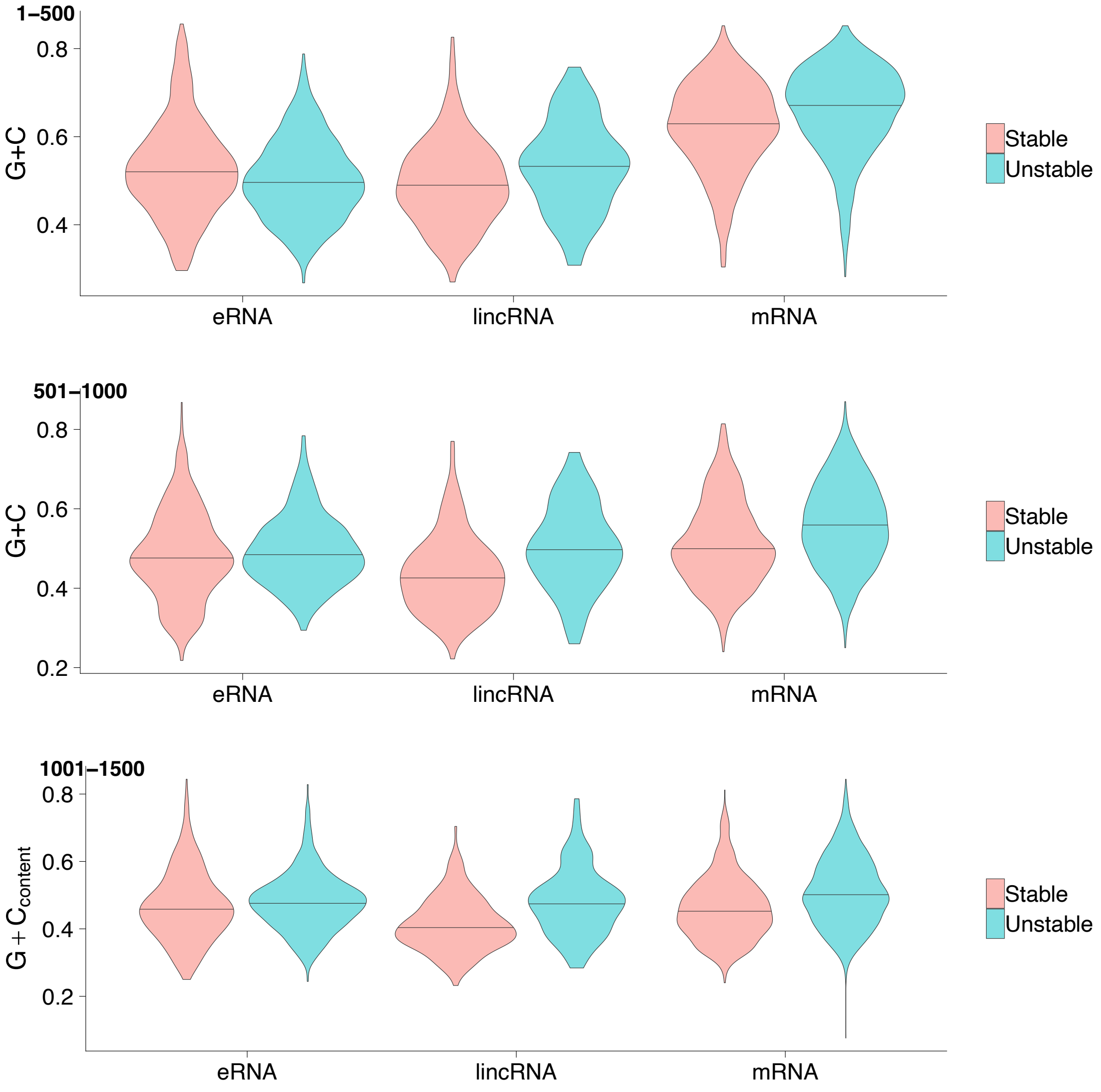


**Supplemental Figure 20**. **G+C content in intervals downstream of the TSS for various classes of transcription units**. Results are shown for intervals 1-500bp (*top*), 501-1000bp (*middle*), and 1001-1500bp (*bottom*) downstream of the TSS.

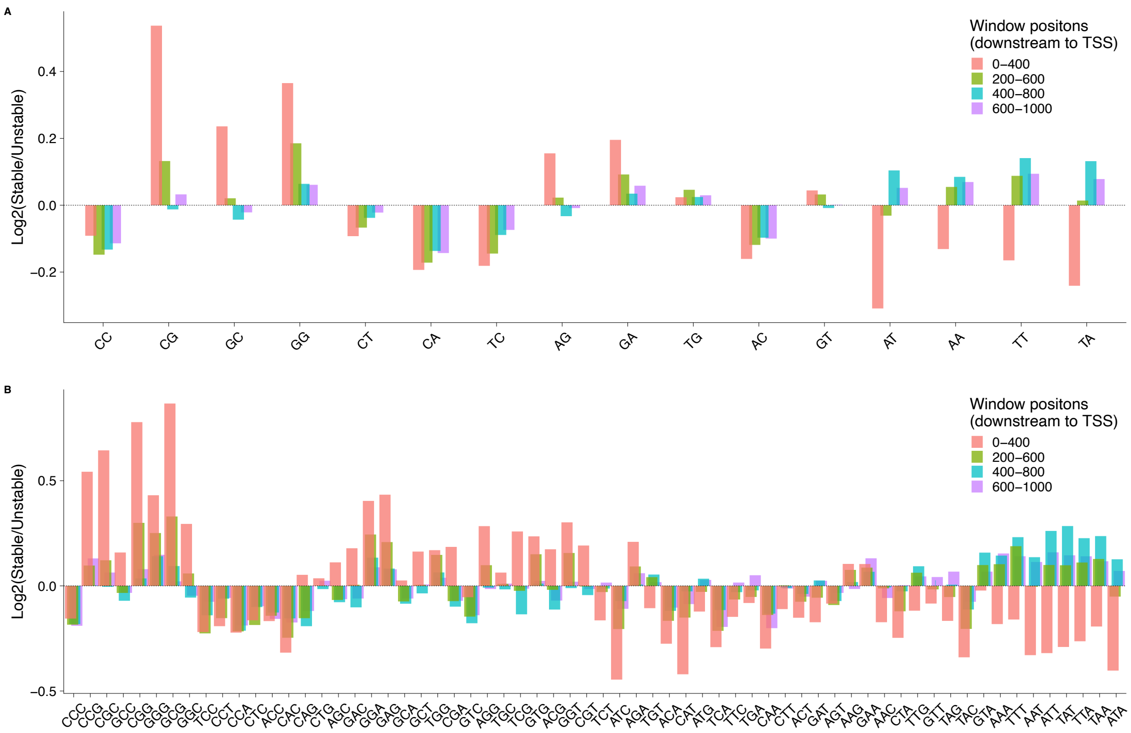


**Supplemental Figure 21. DNA word enrichments in stable transcripts for eRNAs.** For each DNA word, we counted the numbers of total appearances of that word in all stable TUs (taking the top 20% by estimated half-life), $c_{s,w}$, and in all unstable transcripts (the bottom 20%), $c_{u,w}$. We then measured the stability enrichment as $\log_2(c_{s,w}/c_{u,w})$ (see **Methods**). The analysis was repeated in 400bp windows at four different distances downstream of the TSS (see legend). (A) Results for 2-mers. (B) Results for 3-mers. The order of the k-mers reflects the ($c_{s,w}/c_{u,w}$) ratio in the first mRNA window (0-1000bp; see **Supplemental Fig. 18**).

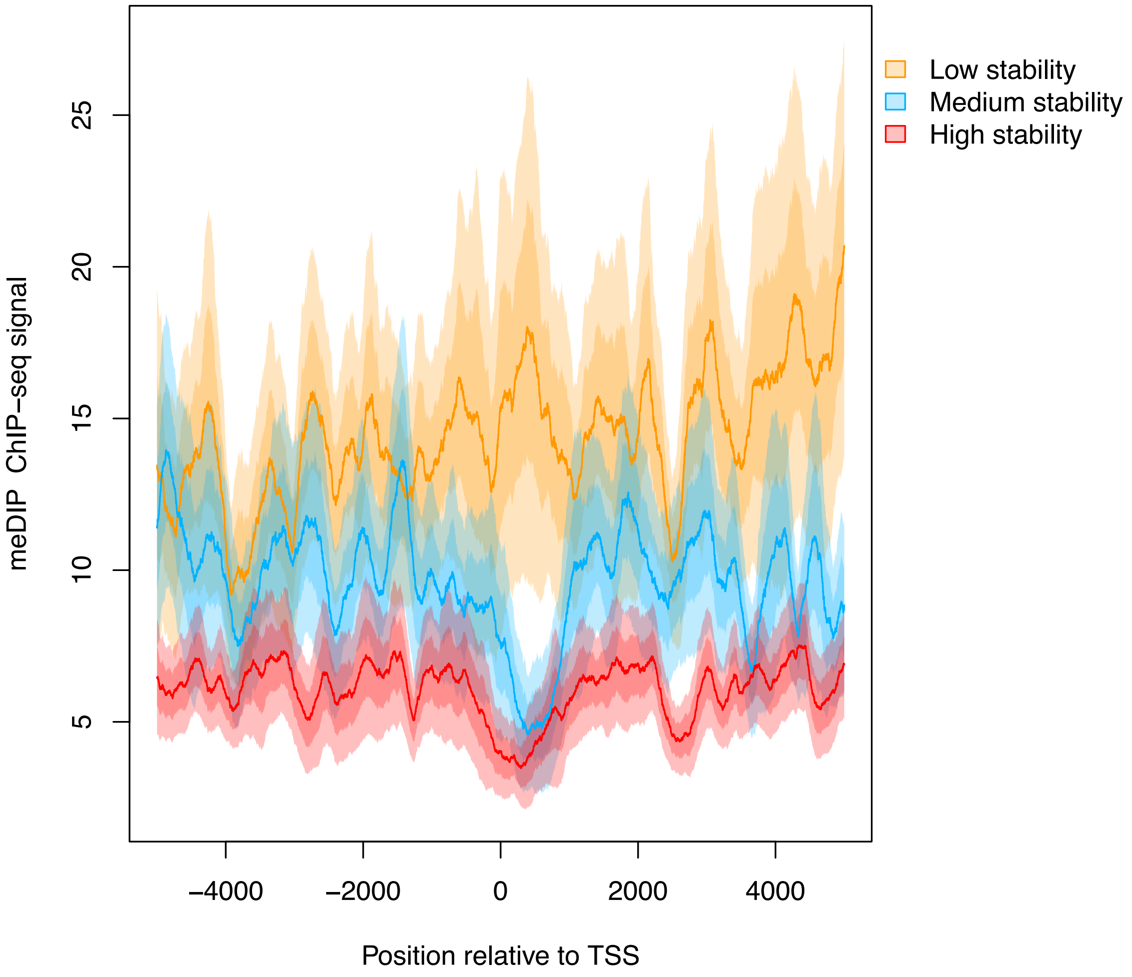


**Supplemental Figure 22. DNA methylation in lincRNAs of various stability levels.** Plots represent the average signal of the methylated DNA immunoprecipitation (MeDIP-seq) assay in K562 cells in three sets of lincRNAs: Low stability (lowest 20% by estimated half-life); Medium stability (40%-60%); and High stability (highest 20%).

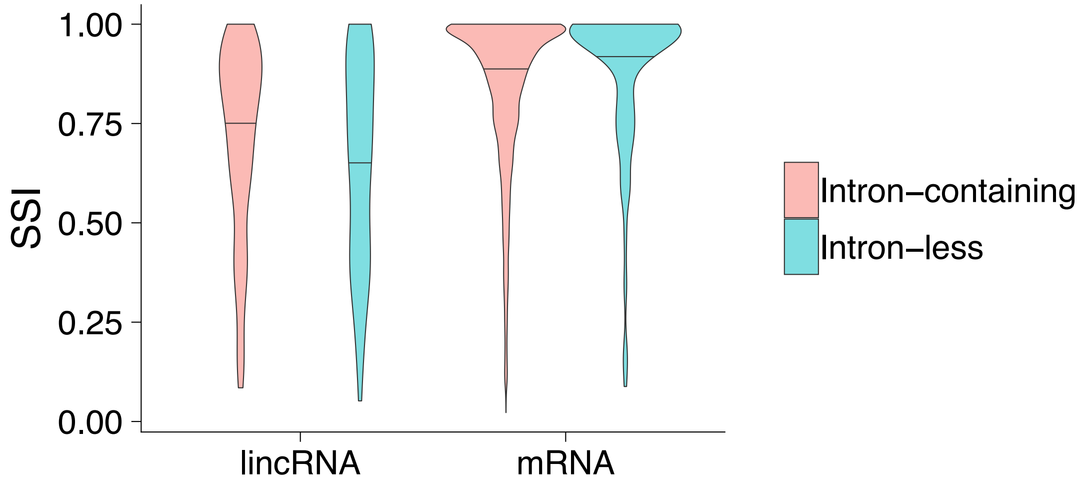


**Supplemental Figure 23. Sequence Stability Index of intron-containing versus intron-less genes:** Results are shown for protein-coding mRNAs and lincRNAs.

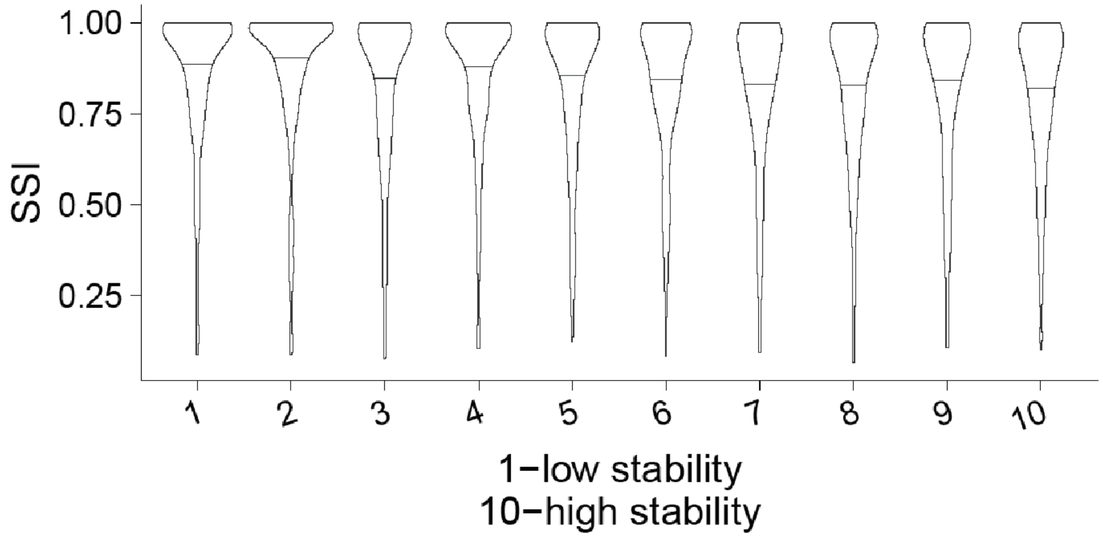


**Supplemental Figure 24. Sequence Stability Index (SSI) for mRNAs of various stability classes.** Each category represents a decile ranging from lowest stability (1) to highest stability (10), based on the estimated half-lives.

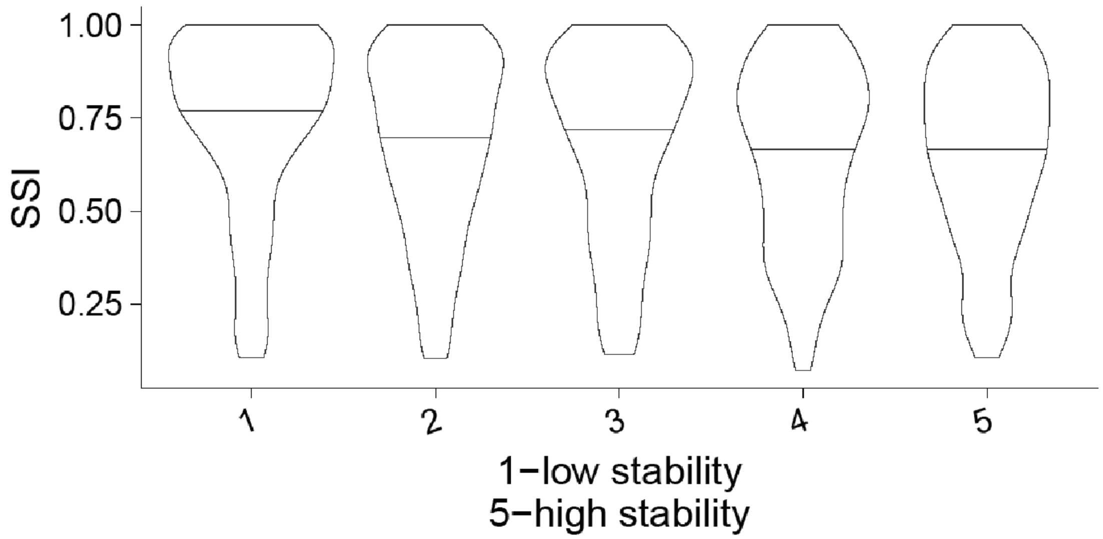


**Supplemental Figure 25. Sequence Stability Index (SSI) for lincRNAs of various stability classes.** Each category represents a quintile ranging from lowest stability (1) to highest stability (5), based on the estimated half-lives.

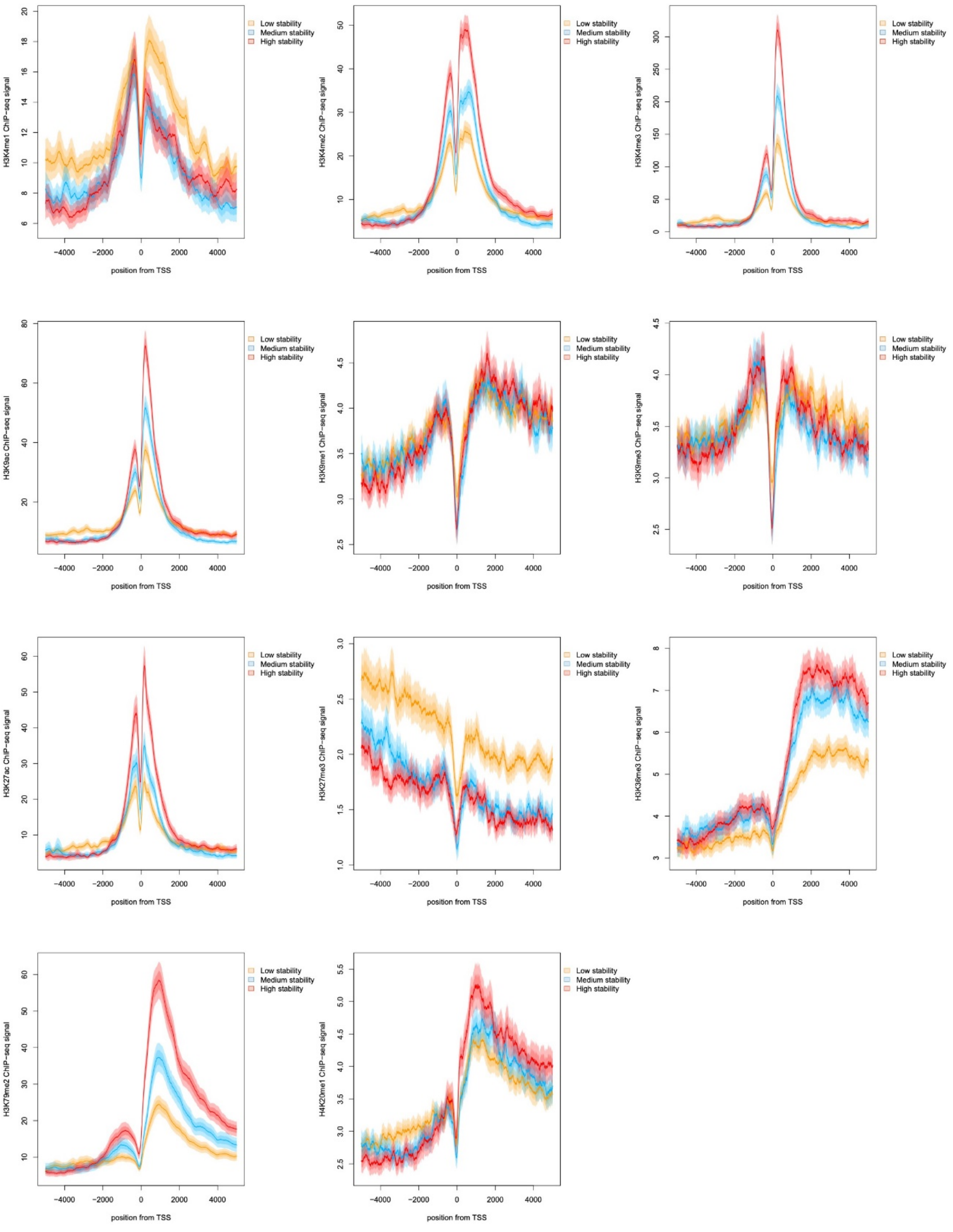


**Supplemental Figure 26**. **Histone modification signals for protein-coding mRNAs of various stability classes.** 11 different histone-modifications are shown. In each panel, separate plots are shown for mRNAs in three stability classes: Low stability (lowest 20% by estimated half-life); Medium stability (40%-60%); and High stability (highest 20%).

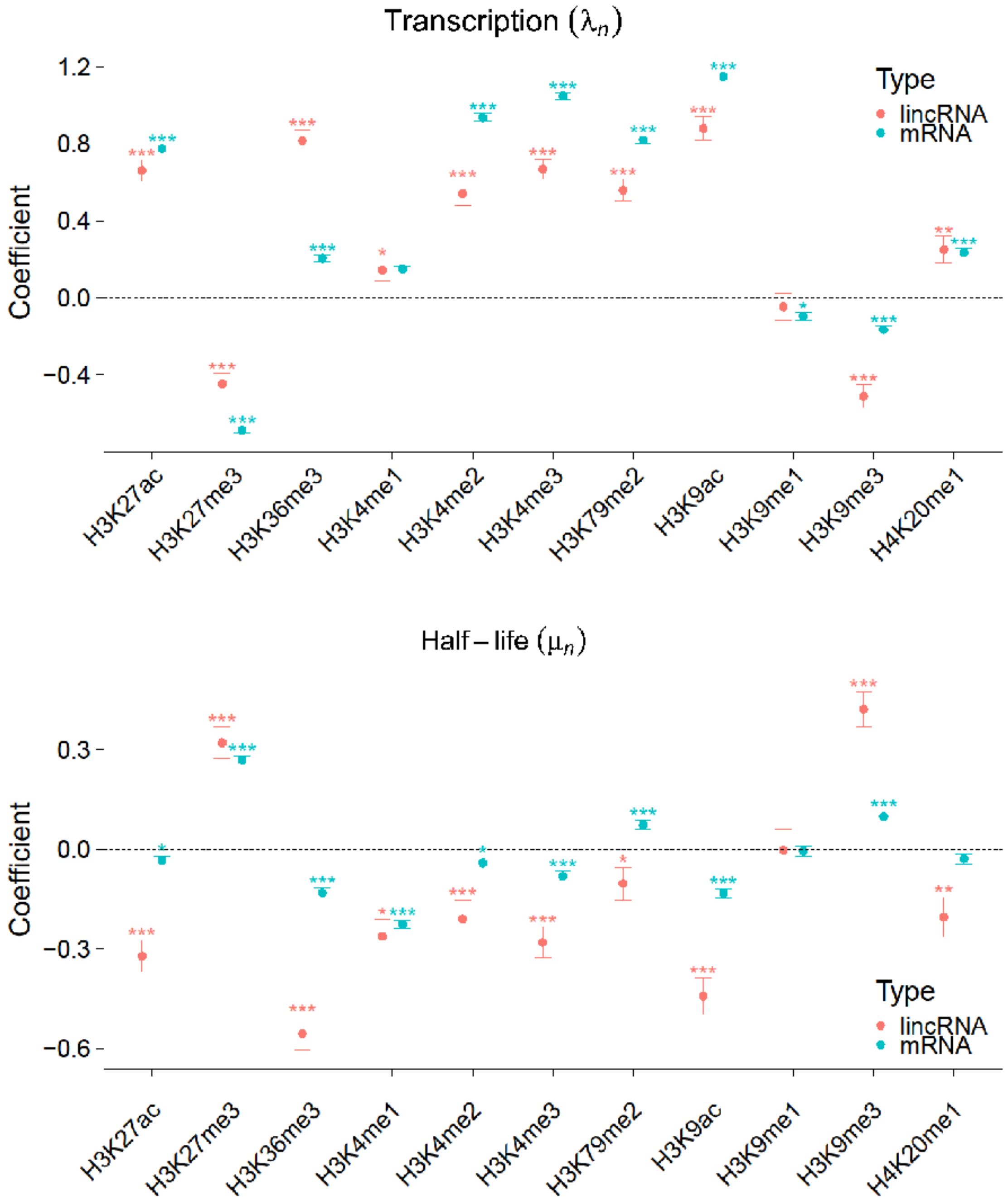


**Supplemental Figure 27.** Estimated SEM coefficients for transcription ($\lambda_n$; top) and half-life ($\mu_n$; bottom) for 11 histone modifications, as assayed by ChIP-seq in the 500 bases immediately downstream of the TSS. The entire set of genes of each type was analyzed. Error bars and significance are as in **Fig. 3B**

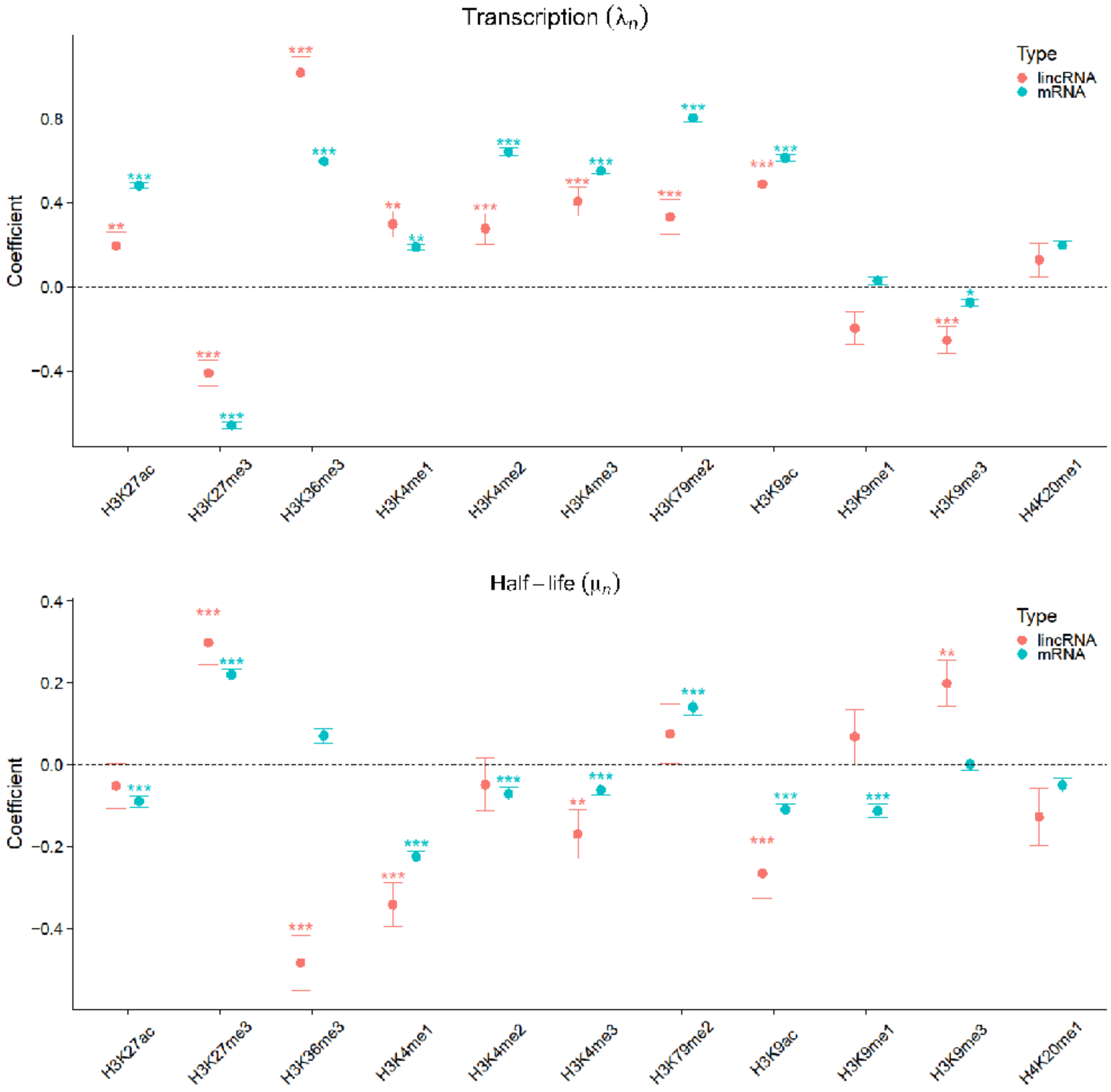


**Supplemental Figure 28.** Estimated SEM coefficients for transcription ($\lambda_n$; top) and half-life ($\mu_n$; bottom) for 11 histone modifications, as assayed by ChIP- in the 1000-1500 bases downstream of the TSS. The entire set of genes of each type was analyzed. Error bars and significance are as in **Fig. 3B**

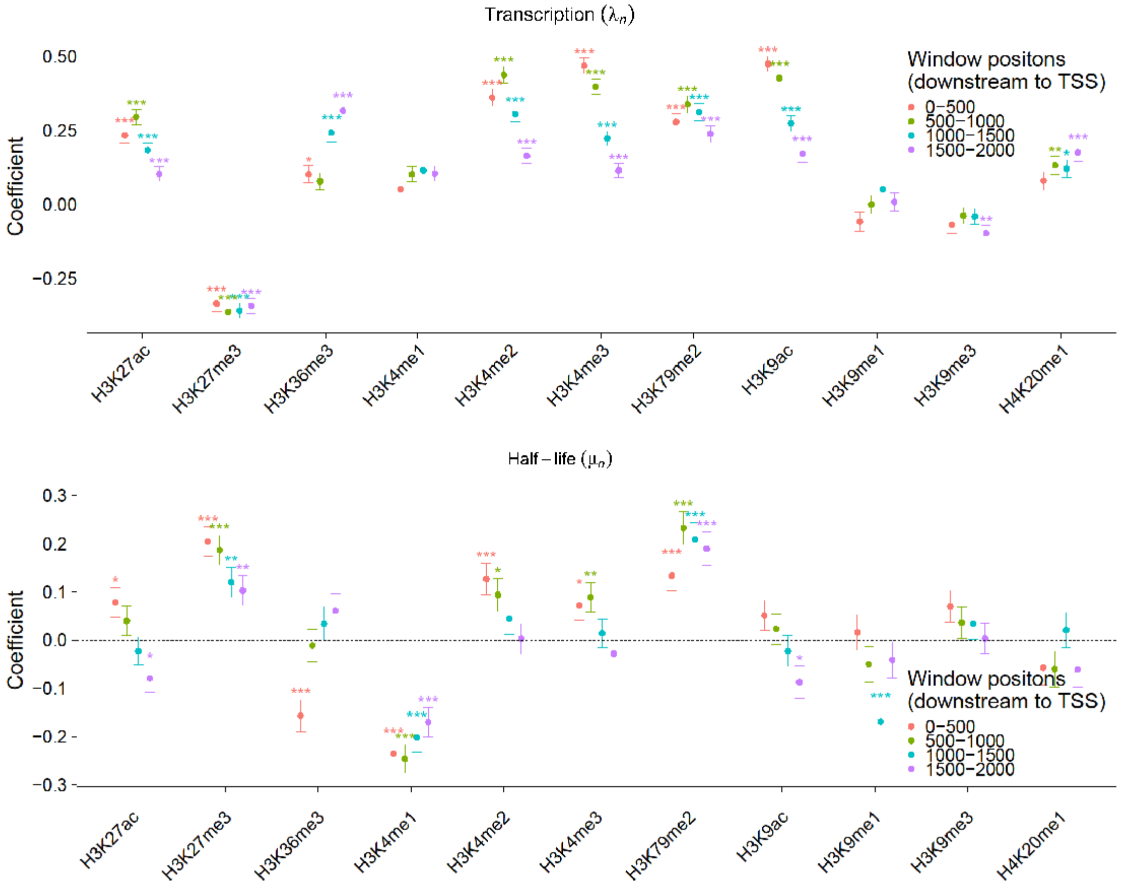


**Supplemental Figure 29.** Estimated SEM coefficients for transcription ($\lambda_n$; top) and half-life ($\mu_n$; bottom) for 11 histone modifications, as assayed by ChIP-seq in 4 windows, each of 500 bases downstream of the TSS (0-500, 500-1000, 1000-1500, 1500-2000, represented by colors). The entire set of mRNA genes was analyzed. Error bars and significance are as in **Fig. 3B**
